# Understanding the roles of secondary shell hotspots in protein-protein complexes

**DOI:** 10.1101/2024.08.26.609822

**Authors:** Parvathy Jayadevan, Yazhini Arangasamy, Narayanaswamy Srinivasan, Ramanathan Sowdhamini

## Abstract

Hotspots are interfacial residues in protein-protein complexes that contribute significantly to complex stability. Methods for identifying interfacial residues in protein-protein complexes are based on two approaches, namely, (a) distance-based methods, which identify residues that form direct interactions with the partner protein and (b) Accessibility Surface Area (ASA)-based methods, which identify those residues which are solvent-exposed in the isolated form of the protein and become buried upon complex formation. In this study, we introduce the concept of secondary shell hotspots, which are hotspots uniquely identified by the distance-based approach, staying buried in both the bound and isolated forms of the protein and yet forming direct interactions with the partner protein. From the analysis of the dataset curated from Docking Benchmark 5.5, comprising of 94 protein-protein complexes, we find that secondary shell hotspots are more evolutionarily conserved and have distinct Chou-Fasman propensities and interaction patterns compared to other hotspots. Finally, we present detailed case studies to show that the interaction network formed by the secondary shell hotspots is crucial for complex stability and activity. Further, they act as potentially allosteric propagators and bridge interfacial and non-interfacial sites in the protein. Their mutations to any other amino acid types cause significant destabilization. Overall, this study sheds light on the uniqueness and importance of secondary shell hotspots in protein-protein complexes.

## 1. INTRODUCTION

Hotspots are interfacial residues in protein-protein complexes that contribute the most to the stability. They are defined as those residues whose mutations to Alanine cause a significant decrease in the binding strength, indicated by a ΔΔG_binding_ greater than 2.0 kcal/mol^1,2^. Hotspots have been shown to overlap with structurally conserved residues^3,4^. It has been proven that the hotspots are usually surrounded by residues that are less conserved and not important for binding, but they shelter the hotspots from the solvent^4,5^. The hotspot generally occurs as discontinuous clusters at the interface known as hot regions. The residues within the same cluster have been shown to exhibit cooperativity among them^4–7^. An earlier study by Bogan et al.^5^ has shown that the hotspots have an amino acid composition distinct from the other interface residues. Trp, Arg, and Tyr are the most frequently found hotspots. The higher propensity of Trp is attributed to its large size, due to which a mutation to Ala creates a large cavity, causing complex destabilization. It can form aromatic pi interactions and can also be a H-bond donor. Similarly, Tyr can form stabilizing aromatic pi interactions and H-bonds using its hydroxyl group, and Arg can form up to five H-bonds and a salt bridge. Leu, Ser, Thr, and Val residues are disfavored as hotspots^5^. Impaired PPIs can cause many diseases, such as neurological disorders and cancer. Moreover, the conserved structure of hotspots and their tremendous impact on the binding energy have made them attractive medical targets for designing inhibitor drugs^8^. Such inhibitors can avoid unwanted protein-protein interactions and more effectively treat various diseases^1,5,9,10^. Such drugs are designed by targeting hotspots with virtual ligand screening and template-directed combinatorial chemistry^11^. In addition, prior knowledge of hotspot residues has been extensively employed in protein−protein docking^11–14^.

Experimentally, hotspots are most commonly identified using Alanine scanning mutagenesis, where residues in a protein of interest are mutated to Ala, one at a time, to then study its consequence on protein stability and/or activity^15^. The Ala substitutions remove sidechain atoms past the β-carbon without introducing additional conformational freedom. Thus, the role of sidechain functional groups at specific positions and the energetic contributions of individual sidechains to protein binding can be inferred from Ala mutations. Other experimental methods for hotspot identification include Ala shaving and residue grafting^16^. The alanine scanning results are compiled in a database (Alanine Scanning Energetics Database, ASEdb)^2^. The Binding Interface Database (BID)^17^ contains the experimentally verified hotspots. However, these methods are time-consuming and labour-intensive as they require the purification and analysis of the mutant proteins. Hence, to complement these experimental techniques, several fast and accurate computational methods have been developed over the years^1,9,13,18–20^. FoldX^21^ and Robetta^22^ predict hotspots through computational alanine scanning and calculate the associated change in binding energy. Several empirical methods are also employed in hotspot prediction^23–26^. For example, the PPCheck^23^ web-server computes the energy values corresponding to the non-bonded interactions between the proteins in the complex, namely hydrogen bonding, van der Waals (vdW), and electrostatic interactions. The sum of these values is normalized by the number of interactions the residue makes, and this normalized energy is further used for hotspot predictions. ECMIS^24^ is another empirical method that uses a sum of energy score, conservation score, and mass index score to predict hotspots. Several machine learning-based algorithms have been developed to predict hotspots^27–34^. Few are solely based on sequence information^27,28,35^. Nguyen et al. proposed a new set of primary sequence-based descriptors derived using *in silico* alanine scanning and digital signal processing techniques that could distinguish hotspots from other interface residues. These descriptors were further utilized in a machine learning model with a random forest classifier, which has an accuracy of 79%^28^. Higa and Tozzi developed a tool based on structural features (like amino acid type and surface area) and evolution-related features (like conservation score) with a support vector machine (SVM) algorithm to predict hotspots given only the unbound structures of the constituent proteins^32^. HSPred is a combination of machine learning and energy-based methods that use SVM to predict hotspots, given the structure of the complex^33,34^. In this method, the energy terms that contribute to hotspot interactions, i.e. vdW potentials, solvation energy, hydrogen bonds and Coulomb electrostatics, are used as features to develop a machine learning model. KFC2 is a server based on various structural features like flexibilities, atomic densities, and ASA that uses the SVM algorithm^30,31^.

Methods for identifying interfacial residues in protein-protein complexes are (a) distance-based, which identifies residues that form direct interactions with the partner protein, or (b) Accessibility Surface Area (ASA)-based, which identifies those residues which are solvent-exposed in the isolated form of the protein and become buried upon complex formation. In an earlier work^36^, we observed that the hotspots unique to the distance-based method stay buried in both the isolated and bound forms of the protein, compared to the other two categories of hotspots (namely, hotspots unique to ASA and hotspots common to ASA and distance). Hence, we coined the term “secondary shell hotspots” for the hotspots that are unique to the distance method since they lie beneath the primary layer of hotspots and are less accessible to the solvent. The current study analyzes the secondary shell hotspots in the 94 protein-protein complexes newly curated from the Docking Benchmark 5.5^37,38^. We observe that the evolutionary conservation scores, Chou-Fasman propensity, and interaction pattern show contrasting trends for secondary shell hotspots compared to the other two categories of hotspots. Besides, the detailed case study analyses of five complexes further highlighted the unique roles of secondary shell hotspots.

## 2. MATERIALS AND METHODS

### 2.1 Dataset preparation

The dataset used in this study was curated from the protein-protein Docking Benchmark 5.5^37^, which contains experimental structures of 257 protein-protein complexes from the Protein Data Bank (PDB)^39^. Docking Benchmark 5.5 is widely used since it contains the unbound structures of the constituent proteins for each protein-protein complex. All the structures in this dataset have a resolution better than 3.25 Å, amino acid length greater than or equal to 30 amino acids, and no missing residues at the interface. Additionally, the complexes in the dataset are non-redundant at the SCOPe family level^40^. Besides these criteria implemented in Docking Benchmark 5.5, entries that were non-dimers in the biological assembly and those with missing residues were removed. The resulting dataset consists of 129 protein-protein complexes, of which only 94 contain secondary shell hotspots (Figure S1A). This final dataset of 94 structures consists of diverse categories of protein-protein complexes, namely, (a) enzyme-containing, (b) antibody-containing, (c) G-protein-containing, (d) receptor-containing, and (e) Others, miscellaneous (48.9%, 9.6%, 10.6%, 8.5%, and 22.3%, respectively) (Figure S1B).

### 2.2 Identification of secondary shell hotspots

The methods used for identifying interfacial residues in protein-protein complexes are generally based on one of the two following approaches: (a) distance-based method^41–43^, which identifies residues that form direct interactions with the partner protein based on a chosen distance cut-off between residue atoms, and (b) ASA-based method^44,45^ which identifies residues that are solvent-exposed in the isolated form of the protein and become buried upon binding with the partner protein, as interface. In an earlier work^36^, we observed that even though these two approaches are widely used, their predictions differ. In fact, the interface residues uniquely identified by these two approaches vary significantly in terms of their properties like conservation pattern, contribution towards overall binding energy, etc., as well as in terms of their roles in complex formation, stability, and dynamics^36^. Hotspots are interface residues that contribute substantially to the stability of the protein-protein complex to the extent that their mutations lead to destabilization or sometimes disruption of the whole complex. A hotspot can be uniquely identified by the distance-based method but failed to be captured as an interface residue by the ASA-based method due to one of the following three scenarios: (1) it stays buried in both the isolated and bound forms of the protein, (2) it is buried in the isolated form and solvent-exposed in the bound form or (3) it stays solvent-exposed in both the isolated and bound forms of the protein. It was intriguing to observe that most of the hotspots uniquely identified by the distance-based approach fall under the category (1), i.e., they stay buried in both the isolated and bound forms of the protein^36^. Hence, we termed such hotspots unique to the distance method as “secondary shell hotspots” as they lie beneath the primary layer of hotspots at the interface. We verified if this observation is specific to the dataset chosen in the earlier study. Hence, we first identified the interfacial residues using the distance-based and ASA-based criteria on the dataset curated as described in Section 2.1. The distance-based approach identifies those residue pairs as interacting if any two atoms in the residue are closer than 4.5 Å (Figure S1C). This criterion was implemented using an in-house Python script. The intra-protein interactions in the complexes were identified using the same distance criteria. The interacting pair of residues (both inter-protein and intra-protein) are removed from the analysis if they have short contacts (excluding the short contacts involving Hydrogen atoms) identified using Molprobity^46^. The type of inter-protein and intra-protein interactions were identified using the web-server Proteins Interactions Calculator (PIC)^47^ (in-house) and Residue Interaction Network Generator v3 (RING v3)^48^. The ASA-based approach identifies the centre of the interface as those residues that are well solvent-exposed in the isolated form of the protein, as reflected by the Relative Accessibility Surface Area (RASA) value of greater than 10% and become buried in the complex form with a RASA value less than 7%. Besides, those residues that are partially exposed in the unbound form of the protein (RASA > 7%) and lose the ASA by more than 1 Å^2^ are also part of the interface, as they include residues at the centre as well as the periphery of the interface (Figure S1C). The residues that are placed at the centre of the interface are named “core”, and those at the periphery of the interface are called “rim” residues. Naccess^49^ has been used to calculate residue-wise ASA and RASA values in the complex and isolated forms of the protein. Hotspots are identified *in silico* as those residues in a complex that leads to a ΔΔG > 2 kcal/mol when mutated to Ala^1^. The AlaScan module of FoldX^21^ has been used to determine such residues in each dataset complex. All the comparative analyses in this study have been performed on three categories of hotspots, namely: (a) hotspots unique to distance (secondary shell hotspots), (b) hotspots unique to ASA, and (c) hotspots common to both ASA and distance methods. These will be referred to as (a) SSH, (b) ASA-hotspots, and (c) common-hotspots, respectively, in the following sections. Such comparative analyses would help to identify properties unique to SSHs compared to the other hotspots at the interface. Figure 1 shows the PyMol visualization of the Tyr13 of IL4-BP, an SSH in the complex formed by Interleukin-4 (IL-4) and its receptor IL4-BP (PDB code: 1IAR; Figure 1, right panel)^7^, and the same residue in the isolated form of IL4-BP (Figure 1, left panel). Evidently, this residue stays buried both in the bound and isolated forms of IL4-BP, indicated by its RASA values of 1.5% and 1.3%, respectively. Interestingly, more than 90% of the SSH in the current dataset stays buried in both the bound and isolated forms of the protein, which motivated us to analyze this category of hotspots in detail further (Figures S2A).

**Figure 1.**
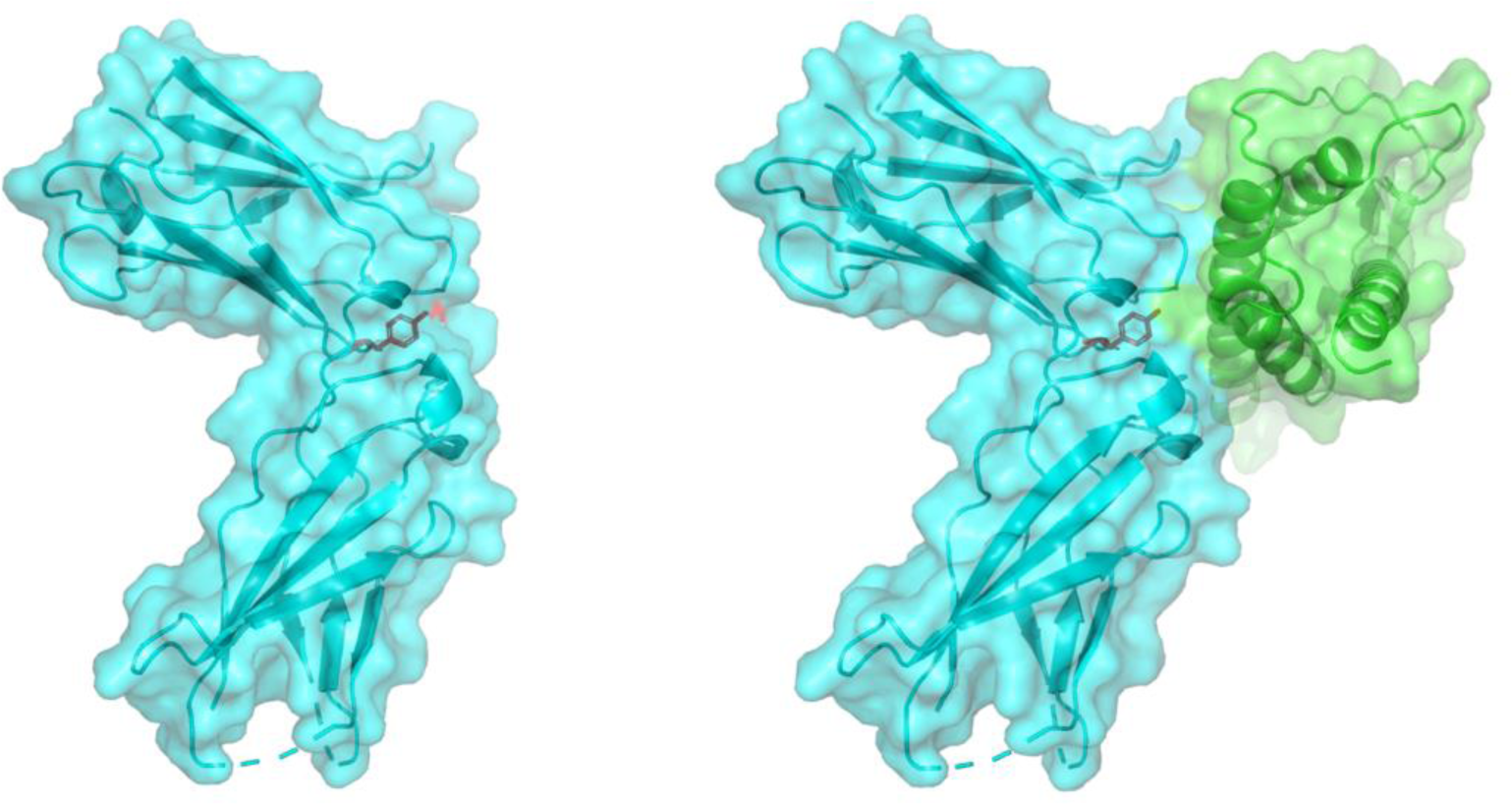
PyMol (The PyMOL Molecular Graphics System, Version 2.5 Schrodinger, LLC) surface view representation of the 1L-4/1L4-BP complex (PDB code: 1IAR) (right), and the isolated form of 1L4-BP (left) with the SSH Tyr13 of IL4-BP shown in red sticks. IL-4 is shown in green, and 1L4-BP in cyan cartoons.

### 2.3 Calculation of residue conservation scores

ConSurf^50^ was used to calculate conservation scores for each of the three categories, as mentioned in Section 2.2. Given an input structure, the ConSurf algorithm identifies homologues using the HMMER method^51^. In this study, we used the Uniprot_TrEMBL database to search for homologues with one iteration and an E-value cut-off of 0.0001. The algorithm finds the representative homologues using the clustering approach implemented by CD-HIT^52^ and uses MAFFT-LINSI^53^ to derive multiple sequence alignments from the identified homologues. From this multiple sequence alignment and the corresponding phylogenetic tree, a Bayesian statistics-based algorithm called Rate4Site^54^ calculates the evolutionary rates. The ConSurf grades obtained in the final output are the estimated evolutionary rates of each residue position normalized to values ranging from 1 to 9. A grade of 9 indicates the highest conservation and, hence, its importance in the structure and/or function of the protein. Generally, a residue with a ConSurf grade greater than six is considered partially conserved or conservatively substituted, and a grade less than or equal to six is considered poorly conserved. Homologues could not be detected for two complexes in the dataset; hence, the conservation analyses were conducted for the dataset, excluding these two entries. The percentage of each category of hotspots having a conservation score from 1 to 9 is calculated as follows:

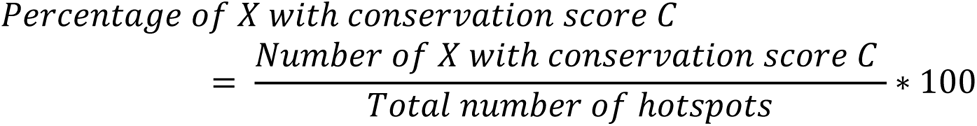

Where X can be SSH, ASA-hotspots, or common-hotspots, C ranges from 1 to 9, and the total number of hotspots is the sum of the number of SSH, ASA-hotspots, and common-hotspots.

### 2.4 Propensity calculations

We employed the Chou–Fasman^55^ propensity calculations to identify the nature of residues with a higher tendency to be an SSH or ASA-hotspots or common-hotspots as follows:

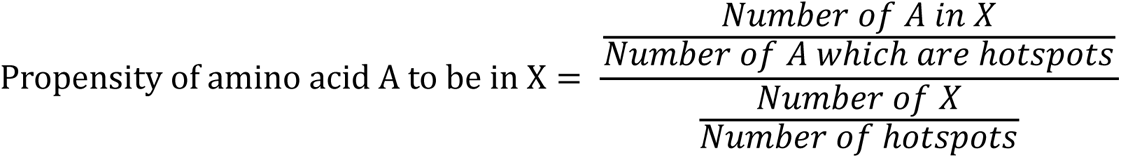

Where X can be SSH, ASA-hotspots, or common-hotspots, and the number of hotspots is the sum of the number of SSH, ASA-hotspots, and common-hotspots.

### 2.5 Prediction of secondary structural elements and Ooi number

To identify if the SSH prefers to be in any specific secondary structural element compared to the other two categories of hotspots, we used software called SSTRUC (D. Smith, unpublished results) for secondary structure assignments. This software implements the Dictionary of Secondary Structure in Proteins (DSSP) algorithm by Kabsch & Sander, which assigns the secondary structures using hydrogen bonding patterns and dihedral angles. The percentage of each category of hotspots belonging to a specific secondary structural element was calculated as follows:

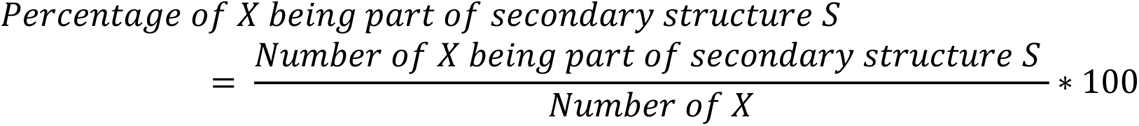

Where X can be SSH or ASA-hotspots, or common-hotspots, and S can be (1) Helix, (2) Beta, (3) Loop, or (4) Ambiguous. Helix includes α, π, and 3_10_ helices; Beta includes β bridge and β bulge, and loop includes turns and coils. Ambiguous indicates a piece of low curvature that cannot be assigned to any hydrogen-bonded structure.

Besides, another parameter, referred to as the Ooi number^56^, was also used from the SSTRUC predictions. The Ooi number of a residue is defined as the number of residues within 8Å from its C^α^ atom. A high Ooi number implies the compactness or high density of the residue’s neighborhood. The percentage of each category of hotspots was calculated as follows:

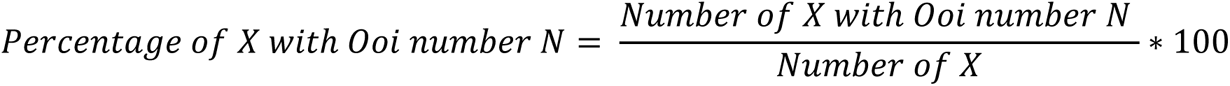

X can be SSH, ASA-hotspots, or common-hotspots, and N can be any integer.

### 2.6 Calculation of depth index

The depth index of an atom is calculated as its distance from the closest solvent-accessible atom. The depth index of a residue is defined as the maximum of the depth index values of all the atoms in the residue. This parameter was calculated using the software Protein Structure and Interaction Analyzer (PSAIA)^57^, which computes the geometric parameters for protein structures, further aiding in investigating protein-protein interaction sites. The current dataset contains the experimental structures for the unbound forms of each protein in the complexes. Hence, we analyzed the change in the depth index of all hotspots upon complex formation, which was calculated as follows:

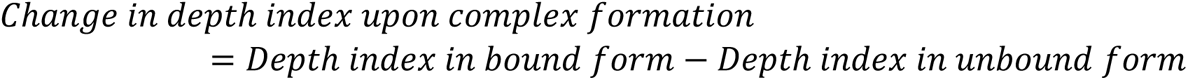

The topologically equivalent residue of each category of hotspots in the bound and unbound structures was identified using TM-align^58^, an algorithm for sequence-independent protein structure comparisons.

### 2.7 Calculation of inter-residue interaction energies

PPCheck^23^ was used to measure the contribution of each category of hotspot residue (SSH, ASA-hotspots, and common-hotspots) to the overall binding energy of the complex. Given a structure with two interacting partners, PPCheck calculates the interchain energies: hydrogen bonding, vdW interaction (hydrophobic interactions), and electrostatics (Figure S4A). The input to PPCheck in this study is the hotspot and its corresponding intra-protein partners in the case of the ASA-hotspots and both inter-protein and intra-protein partners in the case of SSH and common-hotspots so that the local environment of the input residue is preserved^59^. In this analysis, the inter-protein and intra-protein interaction partners were derived based on an all-atom distance cut-off of 10 Å so that none of the interactions predicted by PPCheck are missed.

### 2.8 Perturbation response scanning

Perturbation response scanning (PRS) is a method that evaluates the impact of a single residue perturbation on the entire protein structure. We used the Prody^60^ package to perform PRS on the three case studies to probe into any potential role in allosteric communications unique to SSH. In this method, a protein structure is represented as a mass-spring system where the masses (alternatively, nodes) are C^α^ atoms, and an edge is drawn between two C^α^ atoms if they are within 15 Å. Each residue is perturbed one at a time, at least 1000 times, by exerting a force in random directions and with unit magnitude. The response of each residue to such perturbation is derived as displacements based on Hooke’s law. Residues that cause maximum displacement in the structure upon perturbation are termed effector residues, while the residues that respond maximally to perturbations of all other residues are termed sensor residues. The average value of the effectiveness and sensitivity of all residues in the structure is used as a cut-off to decide effector and sensor residues. Also, to compare effectiveness and sensitivity values across unbound structures and homologous complexes, these values were normalized to values ranging from 0 to 1 using the min-max approach.

### 2.9 Identification of cliques

Protein network models are helpful in elucidating details about the structure, function, and stability of proteins^61–64^. We used GraProStr^61^ to construct the Protein Sidechain Networks (PScN) corresponding to each complex chosen as a case study in which the nodes are residues in the protein structure. The adjacency matrix is constructed by drawing an edge between two nodes only if their interaction strength, I_ij_, is greater than or equal to a chosen cut-off value (I_min_). The adjacency matrix and interaction strength are calculated by GraProStr as follows:

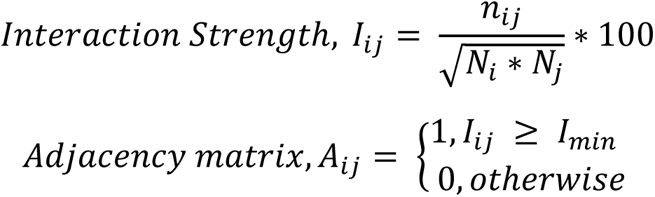

where *n*_*ij*_ is the number of interacting atomic pairs between residues i and j, *N*_*i*_ and *N*_*j*_ are their normalization values and *I*_*min*_ is the interaction strength cut-off (chosen as 1 in this study). From the constructed network, we evaluated a parameter called clique for each case study. A k=n clique is a group of n nodes such that each node is connected to every other node. The default value of k=3 of GraProStr is used in this study. This parameter is used to identify regions of rigidity and higher-order connectivity in protein structures. Besides, interfacial cliques are proven to be useful in analyzing the changes in interactions between subunits for multimeric proteins^64^.

## 3. RESULTS AND DISCUSSION

### 3.1 SSHs are more conserved than other hotspots

In the final curated dataset, there were 4857 interfacial residues, out of which 346 are unique to the distance method, 686 are unique to the ASA method, and 3925 are common to both methods. In this dataset, we computationally identified 1239 hotspots, which include 204 SSH, 80 ASA-hotspots, and 955 common-hotspots. Expectedly, more than 95% of the SSH in the dataset are buried in the bound and isolated forms of the protein, as indicated by their RASA values (<10%) (Figure S2A). But, this inference does not hold for the ASA-hotspots and common-hotspots (Figures S2B and S2C, respectively). This clearly showed that the property of being buried in both the isolated and bound forms is unique to SSH compared to the other hotspots, irrespective of the dataset. This motivated us to analyze further to identify more properties unique to SSH.

Interface residues in protein-protein complexes are highly evolutionarily conserved^3,4^. In addition, our earlier study on interfacial residues showed that interfacial residues unique to distance are more conserved than those unique to ASA^36^. Here, we investigated if the conservation patterns differ for the three categories of hotspots. Figure 2A shows the percentage of the three categories of hotspots with different ConSurf^50^ grades ranging from 1 to 9. A higher percentage of SSH have a ConSurf grade of more than six, which is indicative of a high evolutionary conservation (13.6%, 18.6%, and 44.2% with ConSurf grades 7, 8, and 9, respectively). This is not so for the other categories: ASA-hotspots retain 11.7%, 16.9%, and 15.6% with ConSurf grades 7, 8, and 9, respectively) and the common-hotspots retain 14.6%, 13.6%, and 27.3% with ConSurf grades 7, 8, and 9, respectively. Especially, the percentage of SSH with ConSurf grade 9 is much higher (44.2%) compared to ASA-hotspots (15.6%) and common-hotspots (27.3%), indicating a drastic change in the conservation pattern of SSH in comparison with other hotspots. There has been evidence showing that hotspots, in general, overlap with highly conserved residues at the interface. However, this analysis indicates that SSH can be distinguished from other hotspots, as they are highly evolutionarily conserved.

**Figure 2.**
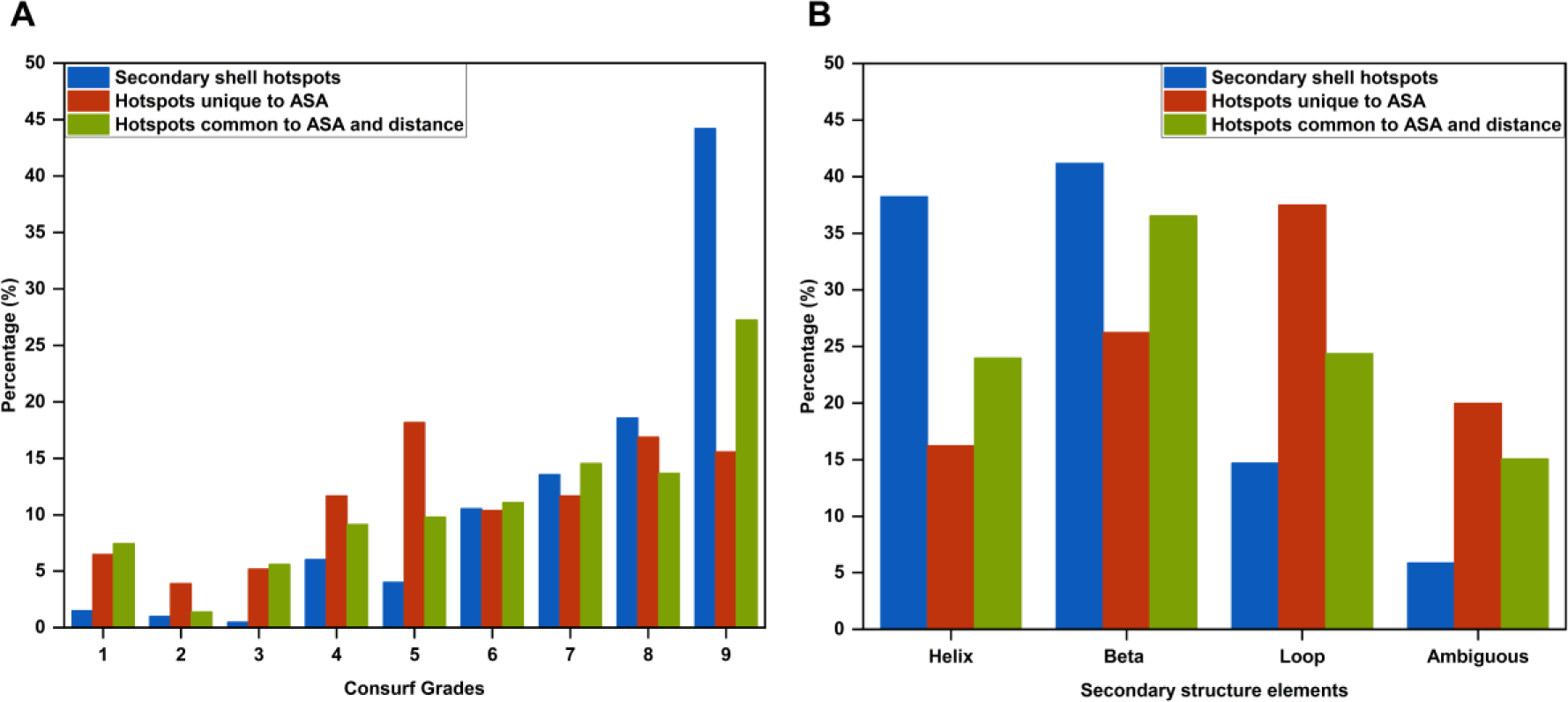
(A) Percentage of SSH, ASA-hotspots, and common-hotspots with ConSurf^50^ grades ranging from 1 to 9. (B) Percentage of SSH, ASA-hotspots, and common-hotspots that are part of secondary structures “Helix”, “Beta”, and “Loops”, as predicted by SSTRUC (D. Smith, unpublished results). “Ambiguous” represents the regions that cannot be assigned to any secondary structure by the DSSP algorithm because of low curvature.

### 3.2 SSHs prefer to be in stable secondary structures

The preference of a residue for different secondary structures reflects its conformational flexibility and potential role in the stability, structure, and function of the protein. Here, we investigate if SSHs have any preference for specific secondary structural elements. Figure 2B shows the percentage of each category of hotspots that are a part of secondary structures, namely helix, beta, and loops. Most SSH prefer to be in helices or sheets (38.2% and 41.2%, respectively) compared to loops (14.7%). 5.9% of them are ambiguous, indicating that they are part of low curvature regions in the protein and that no secondary structure could be assigned using the DSSP algorithm. This result shows that the SSHs tend to be part of stable regions in the protein with restricted conformational flexibility. Compared to SSH, ASA-hotspots, and common-hotspots have a lower preference to be in helices and sheets (16.3%, 26.3% for ASA-hotspots, and 23.9%, 36.5% for common-hotspots, respectively). Notably, ASA-hotspots have the highest percentage of being in loops (37.5%) out of all three categories of hotspots (14.7% and 24.4% for SSH and common-hotspots, respectively) (Figure 2B).

Next, we set out to analyze propensity patterns observed in each hotspot category. For SSH, hydrophobic residues like Val, Leu, Ile, and Met have a propensity greater than one and greater than that of ASA-hotspots and common-hotspots (Figure S3A). This result is anticipated as interface residues are part of hydrophobic patches, and specifically, SSH resides in the hydrophobic core of the protein. Also, for the long-chain residues like His and Lys, the propensities are greater than one and greater than that of ASA-hotspots and common-hotspots. Such long sidechains allow SSH to stay buried and still make interactions with the partner protein. Although Ser shows the highest propensity for SSH, this may be attributed to the low occurrence of Ser hotspots in the entire dataset. Earlier studies have shown that Ser is a less preferred hotspot in general^5^. There are also other polar residues (Thr, Cys, Asn, and Asp), and the aromatic residue Phe with a propensity greater than one and greater than that of ASA-hotspots and common-hotspots. Furthermore, the residues Val, Leu, Ile, Met, His, Lys, Thr, Cys, and Phe, which show a higher propensity to be SSH, are also residues with a higher propensity to be in helices and sheets. This aligns with the observation that a higher percentage of SSH prefer to be in structured secondary structures (Figure 2B). In the case of ASA-hotspot, only Gly, Glu, and Arg have a propensity greater than one (Figure S3A). A higher propensity of Gly agrees with the observation that a higher percentage of ASA-hotspots prefer to be in loops (Figure 2B). The propensity values of common-hotspots corresponding to each of the 20 residue types are comparable (Figure S3A). Overall, this analysis shows that propensity values show contrasting trends for SSH compared to other hotspot categories.

### 3.3 The interaction network and the local environment of SSH are distinct from other hotspots

Since SSH stay buried in the protein core, we anticipated uniqueness for them concerning their interaction patterns and neighbourhood. Figure 3A shows the pair preference for the interactions involving SSH. For this, we first calculated the number of interactions formed by SSH with each of the 20 amino acid types, resulting in a 20×20 matrix. Each row in this matrix corresponds to each amino acid type that belongs to the SSH category. Each element in the row was normalized by the number of interactions made by the respective amino acid type SSH.

**Figure 3.**
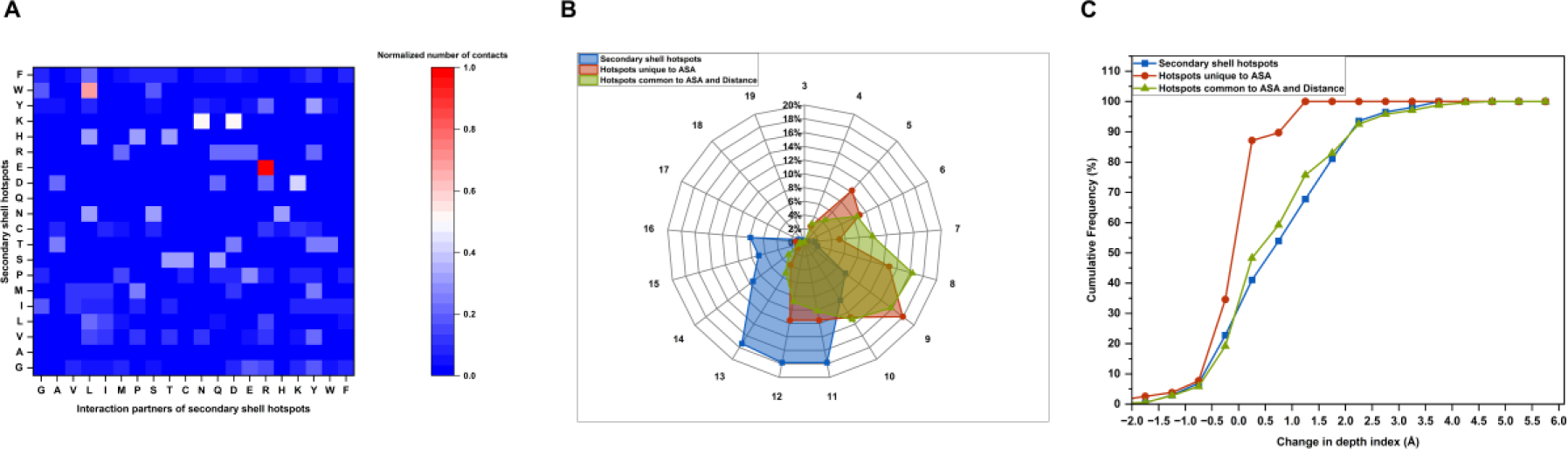
(A) 2D-heat map showing the contact pair preference for each amino acid type that is a SSH. The normalized number of contacts made by each SSH of a specific amino acid type is plotted. The value ranges from 0 to 1, indicated by blue and red, respectively. (B) Radar plot showing the percentage of SSH, ASA-hotspots, and common­hotspots with Ooi numbers ranging from 3 to 19. The Ooi^56^ number for each hotspot is predicted using SSTRUC (D. Smith, unpublished results). (C) Cumulative frequency in percentage of SSH, ASA-hotspots, and common-hotspots with change in depth index upon complexation in bins of size 0.5 A ranging from −2 A Io 6 A. The depth index is computed using PSAIA^57^.

The Glu-Arg pair was the most preferred, with a value of 1, implying that whenever the SSH is a Glu, it interacts with an Arg residue. Similarly, the interaction pairs Lys-Asn, Lys-Asp, and Trp-Leu also have values greater than or equal to 0.5 (0.5, 0.5, and 0.7, respectively). This observation aligns with the fact that hydrophobic and long-chain residues are preferred in SSH, as seen in the Chou-Fasman propensity analysis (Figure S3A).

Next, we analyzed “Ooi number“^56^, predicted by the SSTRUC program. Figure 3B shows a radar plot of the percentage of each hotspot category with various Ooi numbers. Clearly, the three categories have different trends with respect to this parameter. In the dataset, the Ooi numbers varied from 3 to 19. The highest percentage of SSH was observed for Ooi numbers 11, 12, and 13 (17.8%, 17.8%, and 17.3%, respectively). In contrast, the percentages corresponding to ASA-hotspots and common-hotspots for these Ooi numbers are lower compared to the SSH (11.5%, 11.5%, and 3.9% for ASA-hotspots, and 10.18%, 8.78%, and 5.2% for common-hotspots corresponding to Ooi numbers 11, 12, and 13, respectively). For the highest three Ooi numbers, 17, 18, and 19, all three hotspot categories have percentages less than 1. In the following highest Ooi number, which is 16, SSH has the highest percentage comparatively (7.9%, 1.3%, and 0.7% for SSH, ASA-hotspots, and common-hotspots, respectively) (Figure 3B). Overall, this result suggests that SSH have a very dense and compact microenvironment, which is in line with the fact that they prefer to stay buried in both the bound and isolated forms of the protein. However, the Ooi number does not change much as the protein goes from its unbound form to its bound form for all three categories of hotspots (Figure S3B). A significant percentage of hotspots have changed in Ooi values −1, 0, and 1 (19.3%, 48.5%, and 21.3%, respectively, for SSH, 12.8%, 57.7%, and 16.7%, respectively for ASA-hotspots, and 15.1%, 54.3%, and 16.9%, respectively for common-hotspots) (Figure S3B).

We also analyzed the “depth index” to understand if the placement of the residue with respect to the protein surface in the bound and unbound structures of the constituent protein could possibly differentiate between the three categories of hotspots. The cumulative plot for change in depth index upon complexation, defined as in Section 2.6, with bins ranging from −2 Å and 6 Å and bin size of 0.5 Å, was analyzed for each category of hotspot (Figure 3C). A negative value indicates the depth index has reduced, and a positive value indicates that the depth index has increased upon complex formation. A significant change in the cumulative percentage of different hotspots is seen from the bin ranging between −0.5 Å to 0 Å (22.8% for SSH, 34.6% for ASA-hotspots, and 19.2% for common-hotspots), implying that the depth index of a slightly higher percentage of ASA-hotspots tend to decrease as it goes from the unbound to the bound form of the protein. Also, 77.2% of SSH and 80.8% of common-hotspots have a change in depth index value greater than zero, implying that the depth index of most of these hotspots increases upon complex formation. Apart from the possible shielding effect by the partner protein upon complex formation, this observation can be due to conformational changes, which cause these hotspots to get buried more toward the protein’s interior. When we analyze the percentage of intra-protein interactions in unbound form retained upon complex formation (referred to as “retain percentage”), more than half of each category of the hotspot has a retain percentage equal to 100% (53.9%, 66.7%, and 57.9%, for SSH, ASA-hotspots, and common-hotspots) (Figure S3C). This implies that the intra-protein interaction pattern has not changed much for these hotspots upon complex formation. Besides, a significant percentage of hotspots have change in intra-protein partner count only corresponding to the values −1, 0, and 1 (22.8%, 40.6%, and 22.8% for SSH, 19.2%, 46.2%, and 23.1% for ASA-hotspots, and 18.9%, 47.6%, and 15.8% for common hotspots, respectively) (Figure S3D). Overall, the results suggest that along with retaining the interaction pattern upon binding with the partner protein, there is also no gain in the intra-protein interactions in the bound form. Hence, the observed increase is due to the shielding effect by the partner protein.

### 3.4 Residue-wise energy can distinguish SSH from ASA-hotspots

Hotspots are known to contribute significantly to the overall binding energy. Hence, to understand if SSH has any unique role in contributing towards the interaction energy, we analyzed the residue-wise energy of each hotspot using PPCheck^23^ and studied the cumulative percentage of each category of hotspot in various bins of size 100 kJ/mol ranging from −600 kJ/mol to 800 kJ/mol (Figure S4A-B). The cumulative plot showed that 79.8% of SSH, 29.5% of ASA-hotspots, and 79.6% of common-hotspots have residue-wise energy less than −100 kJ/mol. 99.5% of SSH, 100% of ASA-hotspots, and 98.9% of common-hotspots have residue-wise energy less than 0 kJ/mol, which implies a favourable contribution towards the overall binding energy of the complex. Clearly, there is no significant change beyond the energy value of 0 kJ/mol. Overall, we observe that all hotspots tend to contribute favourably towards the interaction energy of the complex (residue-wise energy less than 0 kJ/mol). However, SSH and common-hotspots tend to contribute more favourably than ASA-hotspots (Figure S4B). This could be due to the stabilizing interactions formed by the SSH and common-hotspots with the partner protein.

### 3.5 Case studies

To investigate any specific roles SSH holds in protein-protein complexes, we performed a detailed analysis of three case studies, as discussed in the following sections.

#### 3.5.1 Case study 1

β-lactamases are enzymes in bacteria that give them resistance to β-lactam antibiotics, for example, penicillins and cephalosporins. They catalyze the hydrolysis of the amide bond in the β-lactam ring to render them inactive. TEM-1 is a class A β-lactamase. *Streptomyces* species secrete proteinaceous inhibitors of class A β-lactamases with exceptionally high affinities. β-lactamase inhibitor protein (BLIP) is one such inhibitor from *Streptomyces clavuligerus* that exhibits Ki values in the nanomolar to picomolar range for various class A β-lactamases like TEM-1. We performed detailed analyses on the experimental structure of BLIP bound to TEM-1, solved at a resolution of 1.7 Å (PDB code: 1JTG) (Figure S5A, left panel)^6^. BLIP consists of a tandem repeat of a 76-residue α/β domain. The two domains form a large concave eight-stranded β-sheet upon which aromatic side chains make extensive hydrophobic contacts with the loop-helix region of TEM-1^6^. BLIP-II is another β-lactamase inhibitor protein secreted by the soil bacterium *Streptomyces exfoliatus* SMF19. The experimentally solved structure of BLIP-II in complex with the TEM-1 is also available at 2.3 Å resolution (PDB code: 1JTD) (Figures S5A middle and right panels)^6^. BLIP-II is a seven-bladed β-propeller with a unique blade motif consisting of only three antiparallel β-strands. The TEM-1 binding surface on BLIP-II is exclusive of side chain atoms from residues at the apical β-turns and interblade loop regions that make up the apical rim. BLIP-II binds primarily via its apical face to the same loop-helix region (residues 98–114) of TEM-1 using predominantly hydrophobic contacts, as in BLIP/TEM-1 complex (PDB code: 1JTG). The spanning of BLIP-II across TEM-1 in the complex makes the active site inaccessible to the substrate, but no direct contacts are made with TEM-1 active site residues. Although BLIP-II does not share the same fold as BLIP and has only a pairwise sequence identity of 13.8%, the two proteins show similarities in their binding to TEM-1 (Figures S5A right panel, and Table S1). Both bind to TEM-1 competitively. However, in contrast to BLIP-II, BLIP directly interacts with TEM-1 active site residues. Two protruding β-hairpin loops on BLIP structurally mimic penicillin and are inserted into the TEM-1 active site. Compared with BLIP, BLIP-II binds to TEM-1 more tightly, with lesser interface residues, showing slower dissociation and faster association^6^.

His41 of BLIP is the only SSH in the TEM-1/BLIP complex (PDB code: 1JTG). The Ala mutant of this residue has a ΔΔG of more than 2 kcal/mol and causes an increase in the Ki value indicative of a reduced inhibition capability^65,66^. This residue is part of one of the patches of BLIP hotspots determined as functional epitopes (residues that result in a more than 10-fold decrease in binding affinity when mutated). His41 has a RASA of 0% and 1.5% in the bound and isolated forms of the protein, respectively, and has a ConSurf grade of 4 and residue-wise energy of −3.2 kJ/mol. The topologically equivalent residue of this SSH in BLIP-II is Val175, which is not an interface residue in the TEM-1/BLIP-II complex (PDB code: 1JTD). This motivated us to analyze further and justify the differences due to SSH in one of these homologous complexes, as well as their potential role in specificity.

#### 3.5.2 Case study 2

Ricin is a ribosome-inactivating plant toxin, and in its mature form, it is a 65 kDa glycoprotein consisting of two subunits, RTA and RTB, joined by a single disulfide bond. RTA is an RNA N-glycosidase which inactivates ribosomes in eukaryotes through the cleavage of a conserved ribosomal RNA element known as the sarcin-ricin loop (SRL)^67^. Earlier experimental studies have identified four spatially distinct clusters (I–IV) of toxin-neutralizing epitopes on the surface of RTA. A9 is a single-domain camelid antibody (VHH) that was proposed to recognize a novel epitope on RTA that straddles clusters I and III. A9 has a K_d_ value of 0.1nM and an IC_50_ value of 750 nM with RTA. Here, we analyzed the crystal structure of A9 bound to RTA solved at a resolution of 2.6 Å (PDB code: 6CWG) (Figure S5B left panel)^68^. A9 assumes a classical immunoglobulin fold consisting of nine β-strands arranged in two β-sheets with CDRs 1–3 on one face of the molecule. This structure revealed extensive antibody contact with RTA’s β-strand h, along with limited engagement with α-helix D and α-helix C. Besides, considerable binding affinity, and, consequently, toxin-neutralizing activity of A9 is mediated by an unusual CDR2 containing five consecutive Gly residues that interact with α-helix B, a known neutralizing hotspot on RTA. Removing a single Gly residue from the penta-glycine stretch in CDR2 reduced A9’s binding affinity by 10-fold and eliminated toxin-neutralizing activity^68^. A3C8 is another toxin-neutralizing single-domain antibody, and the crystal structure of A3C8 bound to RTA1-33/44-198 of ricin has been solved at a resolution of 2.7 Å (PDB code: 5SV3) (Figures S5B middle and right panels)^67^. This complex has a K_d_ value of 1.5 E^-8^ M. A3C8 was found to pack with the helical region (residues 90–110) of RTA. A9 and A3C8 have the same SCOP ID (b.1.1.1) and have a pairwise sequence identity of 64.9% and RMSD of 1.8 Å corresponding to the aligned regions after superposition using TM-align^58^, indicating these complexes are homologues (Figure S5B right panel, Table S2).

The ricin/A9 complex (PDB code: 6CWG) has two SSH: 1) Tyr91 of ricin and 2) Cys53 of A9. Cys53 of A9 is reported to form an intra-protein disulfide bridge with A9’s Cys105, which is known to link CDR2 with CDR3^68^. Tyr91 of ricin has an RASA of 3.3% and 5.8%, and Cys53 of A9 has an RASA of 0% and 0.8% in the bound form and isolated forms of the protein, respectively. Tyr91 and Cys53 have ConSurf grades of 8 and 4, residue-wise energies of −6.3 kJ/mol and −4.8 kJ/mol, respectively. The topologically equivalent residues of these SSH in the ricin/A3C8 complex (PDB code: 5SV3) are Tyr91 and Arg53, respectively. Tyr91 of ricin is a SSH in the ricin/A3C8 complex, whereas Arg53 of A3C8 is not.

#### 3.5.3 Case study 3

Interleukin-4 (IL-4) is a crucial determinant for allergy and asthma due to its role as a principal regulatory cytokine during an immune response. Dysregulation of IL-4 function contributes to type 1 hypersensitivity reactions, such as allergies and asthma. IL-4 receptor complex, therefore, represents a promising target for immunotherapy of allergic diseases. IL-4 binds with high affinity (K_d_ ∼ 150 pM) and specificity to the ectodomain of its receptor α chain, IL4-BP. This is followed by the common γ chain (γc) recruitment, which initiates the transmembrane signalling. The ∼140-kDa human IL-4Rα chain has 207 residues in the extracellular domain, 24 in the transmembrane region, and 569 in the intracellular domain^69^. The crystal structure of the complex of IL-4 and its receptor IL4-BP is available at 2.3 Å resolution (PDB code: 1IAR) (Figure S5C)^7^. Even though this complex is not a part of our dataset, we chose this structure as it has therapeutic potential and has a lot of experimental work conducted. IL4-BP has an overall L shape compoesed of two covalently linked domains, D1 (residues 1–91) and D2 (residues 97–197), and folds into a sandwich comprising seven antiparallel β sheets arranged in three-strand and four-strand β-pleated sheets that are twisted against each other by approximately 40⁰. In the complex, IL4-BP binds to the helix AC face of IL-4 and has an almost perpendicular orientation of the L-shaped IL-4BP to the helical axes of αC and αA of IL-4. The IL-4/IL4-BP interface mainly comprises loop L2 from IL4-BP interacting with helix B of IL-4, loops L3 and L1 interacting with αC, and loops L5 and L6 interacting with αA. The loop L4 of IL4-BP connects the domains D1 and D2 and has no interactions with IL-4. The binding epitope reveals a mosaic-like assembly of three discrete clusters of trans-interacting residues. Clusters 1 and 2, with the most contribution towards the complex stability, have an inner core of polar groups which is surrounded by hydrophobic side chains^7^.

Tyr13 of 1L4-BP, a residue part of the loop L1 in domain D1, is the only SSH in this complex. It is also experimentally proven to be a hotspot as an Ala mutation resulted in a ΔΔG of 5.2 kcal/mol, indicative of a strongly reduced affinity more than 500-fold lower than wild-type IL4-BP^70^. The minimal chemical alteration introduced in the Phe variant makes it very unlikely that the observed decrease in binding affinity originates from structural alterations. Instead, it is due to modifications in the sidechain involved in binding^70^. Besides, it is a major binding determinant and a functional sidechain in cluster 1. It forms an H-bond with Glu9 of IL-4, the residue central to cluster 1, and an experimentally verified hotspot^7,71^. Also, its sidechain exhibits cooperative effects with the IL4-BP residues Tyr127 and Tyr183 of cluster 1^70^. Tyr13 has a RASA of 1.3% and 1.5% in the bound and isolated forms of the protein, respectively.

#### 3.5.4 SSH forms interaction networks crucial to binding or activity

Since the SSH stays buried in both the bound and isolated forms of the protein and holds very distinct properties from ASA-hotspots and common-hotspots, we were intrigued to analyze their interaction patterns in all three case studies we had chosen. In case study 1 (CS1) of the TEM-1/BLIP complex (PDB code: 1JTG), the SSH His41 of BLIP forms a favourable vdW interaction with Pro107. This residue is part of the loop-helix region of TEM-1 (99-114) that is experimentally shown to be critical for tight binding with BLIP and BLIP-II and for maintaining the structure and function of TEM-1 (Table 1)^6,72^. Pro107 is also a strictly conserved residue among class A β-lactamases (Table 1)^73^. Experimental mutagenesis of this residue resulted in impairment of binding and activity (Table 1)^6,72,74–76^. Besides, this SSH makes several intra-protein interactions with other residues crucial for binding and activity (Table 1). His41 forms intra-protein interaction with residues like Phe36 and Tyr53 of BLIP, which are common-hotspots according to our criteria as well as experimentally verified to be part of BLIP hotspot patch determined as functional epitope (Table 1)^65,66,74,77^. It forms intra-protein interaction with Tyr50, a common-hotspot and a part of one of the two aromatic patches that account for most of the binding affinity (Phe36, His41, Tyr50, and Tyr53)^66^. Besides, Tyr50 is a specificity determinant in that substitutions at this position result in significant alterations in the relative affinity of BLIP for class A β-lactamases^78^. Similarly, in case study 2 (CS2) and case study 3 (CS3), we observe that the interaction network formed by the SSH is crucial for the complexes in many ways (Tables S3-S5). Hence, we infer that the interactions formed by SSH are essential for complex stability and activity.

**Table 1.**
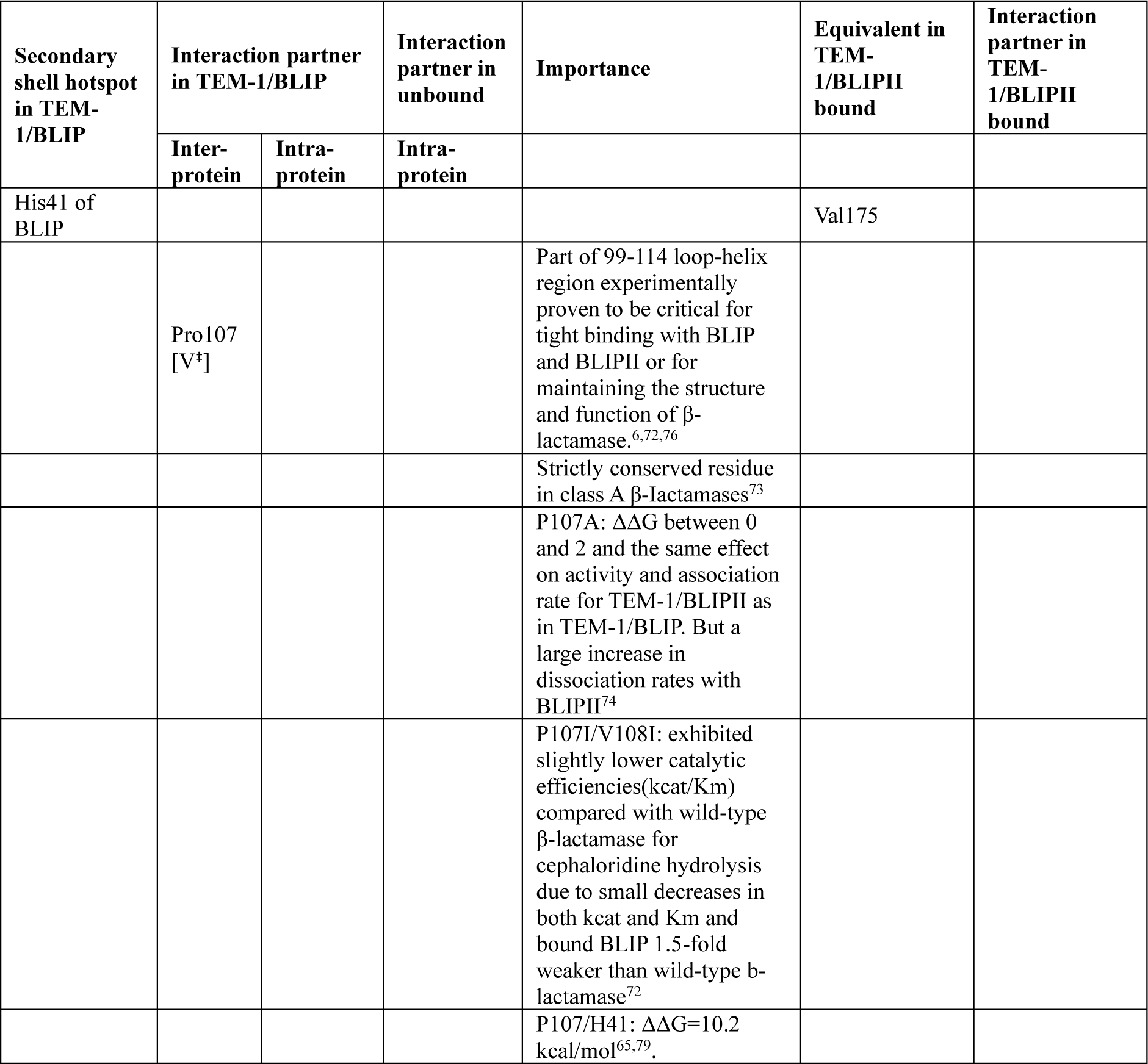
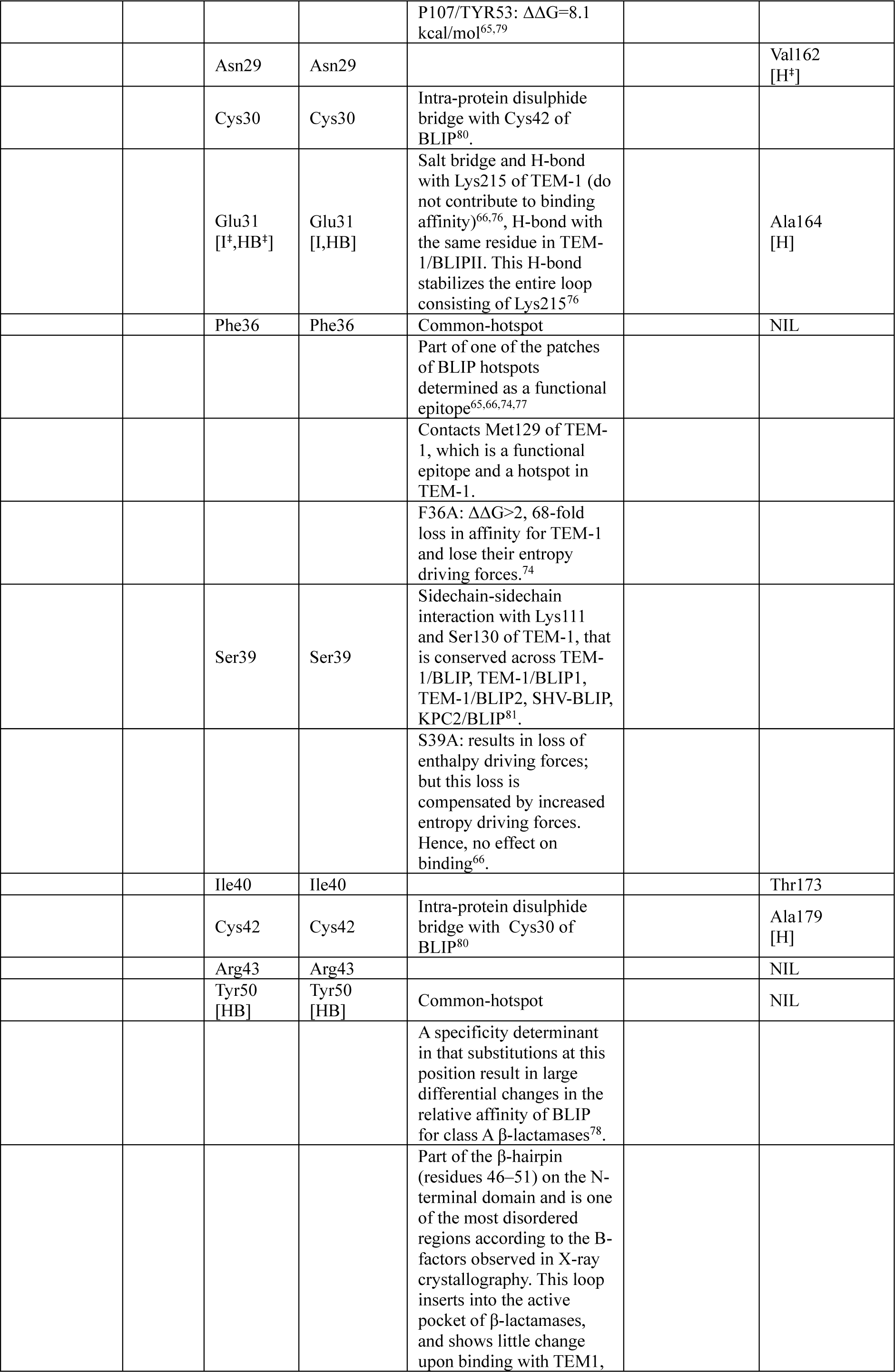
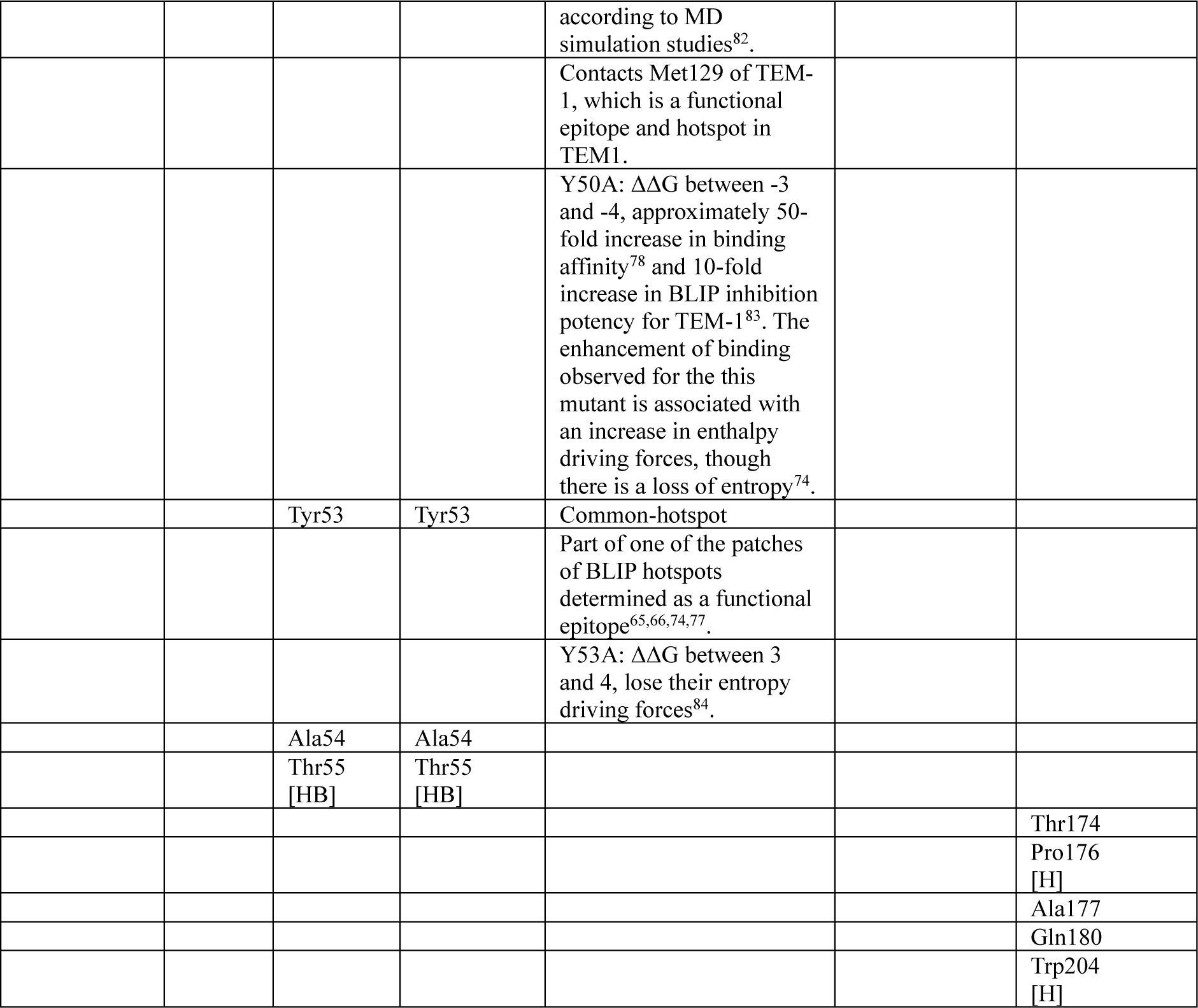
Interaction partners of the secondary shell hotspot His41 of BLIP in TEM-1/BLIP complex (PDB code: 1JTG) of CS1^†^, the unbound structure of BLIP (PDB code: 3GMU), and the equivalent residue Val175 of BLIPII in TEM-1/BLIPII complex (PDB code: 1JTD) of CS1. The inter-protein and intra-protein partners were identified using an all-atom distance cut-off of 4.5 Å, and the nature of interactions was predicted using the PIC^47^ and RING^48^ web-server. H, I, V, and HB represent hydrophobic, ionic, favourable van der Waals, and H-bond interactions, respectively. The importance of the residues in terms of structure, stability, and function was taken from earlier studies.

Since case studies 1 and 2 (PDB codes: 1JTG, 1JTD, and 6CWG, 5SV3) were from the dataset, the unbound structures for the individual proteins in the complex were available. This facilitated the comparison of the SSH’s interaction pattern in the unbound and bound forms of the protein. Evidently, the SSH His41 of BLIP from CS1 (PDB code: 1JTG) has an interaction network that is conserved across the bound and unbound (PDB code: 3GMU) forms of the protein (Table 1). Similarly, the SSH Tyr91 of ricin and Cys53 of A9 from CS2 (Tables S3 and S4, respectively) possess the same characteristics. Hence, we concluded that the SSH conserve their interaction pattern in the unbound form of the protein and complex form with the partner protein. Another interesting observation was that when we compare the SSH Tyr91 of ricin in CS2 (PDB code: 6CWG) and its topologically equivalent residue Tyr91, which is an SSH in the homologous complex ricin/A3C8 (PDB code: 5SV3), the interaction pattern seems to be conserved, including the type of interaction (Table S3). Moreover, Tyr91 in the ricin/A3C8 complex has been proven important for a potent toxin-neutralizing antibody-like A3C8 binding with the helical region of RTA^67^. However, analyzing the other hotspot Cys53 of A9 in the same complex, whose topologically equivalent residue is an Arg residue, which is not an SSH in the ricin/A3C8 complex (PDB code: 5SV3), this property does not hold true. The inter-protein H-bond between Cys53 of A9 and ricin is lost in the case of the topologically equivalent residue Arg53 of A3C8 in the homologous complex (Table S4). Similarly, the intra-protein H-bond that it makes with residues like Gly55, Gly56, and Gly58, which are crucial for binding affinity and toxin-neutralizing activity of ricin/A9 complex, are also lost in the case of Arg53 of A3C8. In CS1, in which the TEM-1/BLIP and TEM-1/BLIPII are not homologous complexes, the interactions formed by the SSH His41 of BLIP are not conserved (Table 1). For example, the favourable vdW interaction that His41 of BLIP makes with Pro107 of TEM-1^6,72^ is absent in the case of Val175, the topologically equivalent residue of His41 in BLIPII. Also, only 4 out of 12 intra-protein interactions made by His41 in the TEM-1/BLIP complex are conserved by Val175 of BLIPII in the TEM-1/BLIPII complex (Table 1). Val175 makes additional intra-protein interactions with Thr174, Pro176, Ala177, Gln180, and Trp204. All observations together shed light on the importance of the interaction network formed by SSH and the potential role of SSH in specificity.

#### 3.5.5 SSHs have a potential role in bridging interfacial and non-interfacial sites in the complex

We performed PRS on all three case studies to understand if SSHs have a role in allostery. Figure 4A shows the average values of effectiveness on perturbing each residue in the two complexes in CS1 and BLIP in the unbound form (PDB code: 3GMU) (Figure 4A left, right and middle panels, respectively). The average effectiveness values of all residues in the complex were chosen as a cut-off above which the residue can be considered an effector residue (0.3, 0.3, and 0.3, respectively for TEM-1/BLIP, TEM-1/BLIPII, and BLIP unbound). It was evident that the SSH His41 of BLIP was an effector residue in TEM-1/BLIP and BLIP unbound form (PDB code: 3GMU), whereas it was not an effector in TEM-1/BLIPII complex. The effectiveness values seem comparable with other interfacial residues in the complex (Figure 4A). Also, the effectiveness value of His41 was reduced from unbound (0.8) to bound (0.4) form of the proteins, and it is different across the TEM-1/BLIPII complex (0.3) (Figure 4B). These observations hold true in the case of the SSH Tyr91 of ricin, and Cys53 of A9 of CS2, and Tyr13 of CS3. The only exception is that Cys53 of A9 is not an effector residue in the ricin/A9 complex but an effector in the ricin/A3C8 complex. This could be due to the salt bridges formed by the topologically equivalent Arg53 in the ricin/A3C8 complex (Figures S6A-B and S7A-B). When we specifically look at the effects caused by the perturbations on SSH, it was observed that such perturbations affect non-interfacial sites in the bound form of the protein. In CS1, the perturbations on the SSH His41 of BLIP cause maximum effect on the residues Asp23, Gly26, Ala27, Glu28, Ala62, and Ala63 of BLIP, which are neither at the interface nor proven to be essential for stability or activity of the complex (Figure 4C left panel). Similarly, in the TEM-1/BLIPII complex (PDB code: 1JTD), perturbations on the equivalent residue Val175 of BLIPII have the highest effect on non-interfacial residues, namely Asp159, Ala177, Glu178, Ser181, and Gly182 of BLIPII (Figure 4C right panel). This also holds true in SSH in CS2 and CS3 (Figures S6C-D and S7C). Notably, in CS3, the perturbations on the SSH Tyr13, which is part of the domain1 of IL4-BP, have the highest effect on residues Ser140, Glu141, Asn142, Asp143, Pro144, Ala145, Asp146, which are part of the domain2 of IL4-BP (Figure S7C).

**Figure 4.**
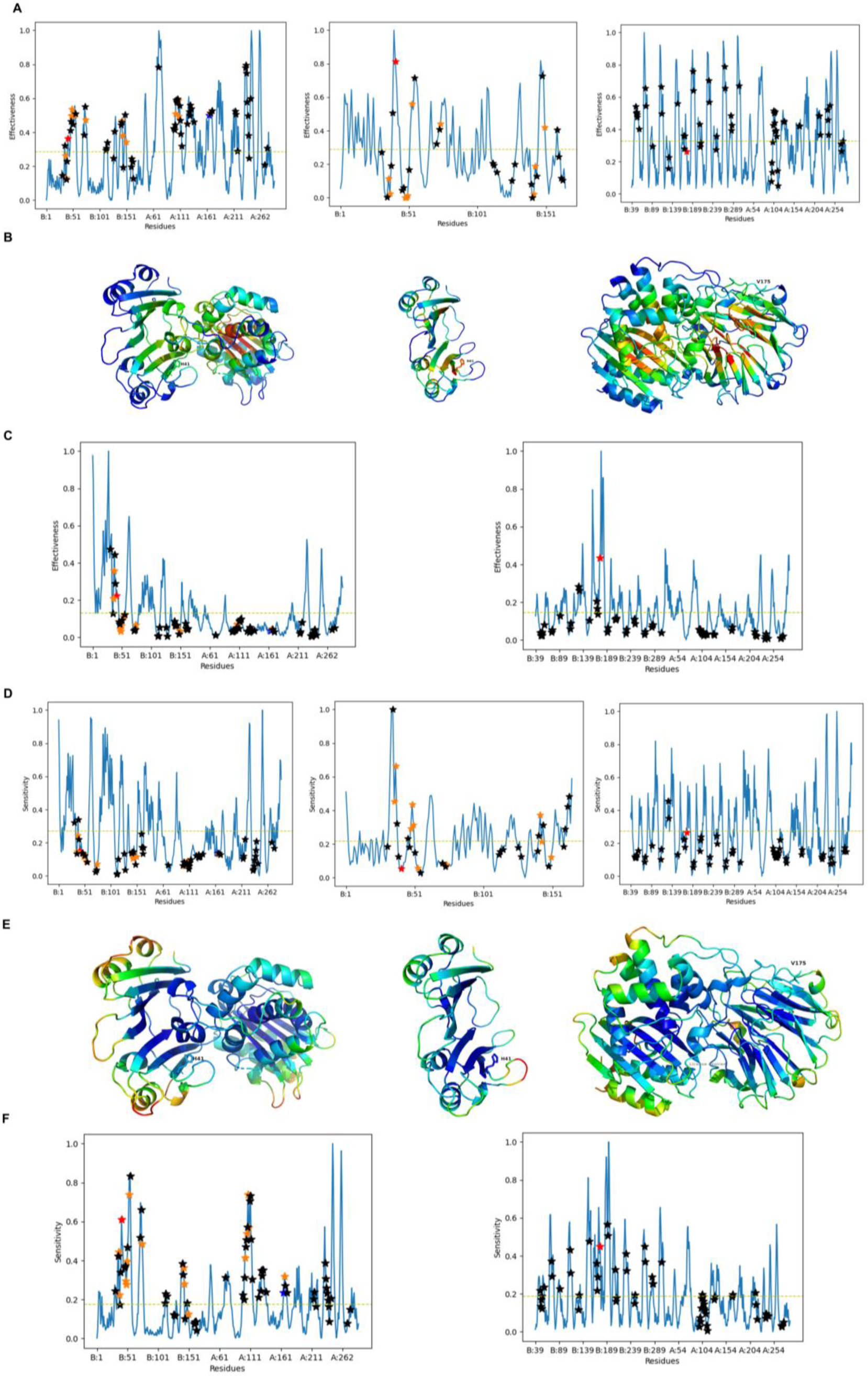
(A) Normalized effectiveness profile and (B) Cartoon representations of TEM-1/BLIP complex (PDB code: 1JTG; left panel), unbound struclure of BLIP (PDB code: 3GMU: middle panel), and TEM-1/BLIPII complex (PDB code: 1JTD: right panel), with effectiveness values mapped (figure generated using PyMol. The PyMOL Molecular Graphics System, Version 2.5 Schrddinger, LLC). (C) Residue-wise effectiveness values for all residues in TEM-1/BLIP (PDB code: 1JTG; left panel) upon perturbing the SSH His41 of BLIP, and for all residues in TEM-1/BLIPII (PDB code: 1JTD: right panel), upon perturbing VaH75 of BLIPII which is topologically equivalent to HIs41 in BLIP, (D) Normalized sensitivity profile and (E) Cartoon representation of TEM-1/BLIP complex (PDB code: 1JTG; left panel), the unbound structure of BLIP (PDB code: 3GMU; middle panel), and TEM-1/BLIPII complex (PDB code: 1JTD; right panel), with sensitivity values mapped (figure generated using PyMol, The PyMOL Molecular Graphics System. Version 2.5 Schrodinger. LLC). (F) Residue-wise sensitivity values of the SSH His41 of BLIP in TEM-1/BLIP complex (PDB code: 1 JTG; left panel) and the equivalent residue Val175 of BLIPII in BLIP In TEM-1 /BLIPII complex (PDB code: 1JTD; right panel), upon perturbing each residue in the complex, These profiles are generated by performing PRS using the Prody package60, The effectiveness and sensitivity values in all profiles are normalized using the min-max method, Mean of effectiveness and sensitivity of all residues Is considered as a cut-off in each profile, above which a residue is considered an effector or sensor (shown as yellow dashed line), The SSH, ASA-hotspots, common­hotspots, and the interfacial residues excluding these hotspots are marked in red, blue, orange, and black stars, respectively.

The sensitivity profiles from PRS analyses of the case studies were also analyzed in detail. From the profile of CS1, it was evident that the SSH is not a sensitive residue in TEM-1/BLIP, TEM-1/BLIPII, and the unbound form of BLIP since the sensitivity was well below the mean values (0.3, 0.3, and 0.2 for TEM-1/BLIP, TEM-1/BLIPII, and the unbound form of BLIP, respectively) (Figure 4D-E). They are the least sensitive in the unbound form of the constituent protein (0.2, 0.3, and 0.1 for TEM-1/BLIP, TEM-1/BLIPII, and the unbound form of BLIP, respectively). A similar trend is also observed in the other case studies (Figures S6E-F and S7D-E). When we analyze the residue-wise sensitivity profiles, we see that SSHs in all three case studies are sensitive to mostly the perturbations on the interfacial residues in the complex (Figures 4F, S6G-H, and S7F). For example, perturbations on the residues Tyr53 (common-hotspot), Thr55 (interface residue common to ASA and distance, but not a hotspot) of BLIP, and Pro107 (common-hotspot) of TEM-1 have a high effect on His41 of BLIP. Tyr53 is a part of one of the patches of BLIP hotspots determined as a functional epitope^65,66,74,77^, and Pro107 is a part of 99-114 loop-helix regions experimentally proven to be critical for tight binding with BLIP and BLIPII or for maintaining the structure and function of β-lactamase (Table 1)^6,72^. Overall, PRS results show that SSH has a potential role as an allosteric propagator bridging interfacial and non-interfacial residues in protein-protein complexes.

#### 3.5.6 SSHs are part of intra-protein and inter-protein cliques

Protein structures can be represented as networks, with the amino acids as nodes and the non-covalent interactions as edges. An analysis of such a network and its associated parameters provides valuable insights into the structure, function, folding, and stability of proteins^61–64^. Hub, clustering coefficient, cluster, cliques, communities, and the shortest paths of communication are some of the most popular network metrics relying on well-established mathematical formulations used to analyze proteins. For example, hubs represent highly connected residues in the protein structure network, which essentially refer to the degree of a node and have been proven to correlate well with experimentally proven hotspots. We constructed weighted sidechain networks of all the case study complexes, as discussed in Section 2.9. In this study, we analyzed a parameter called clique, which indicates regions of rigidity and high-order connectivity in proteins. In CS1, we identified one inter-protein clique involving the SSH His41 of BLIP, which consisted of Pro107 of TEM-1 and Tyr53 of BLIP. These are interaction partners of the SSH and have been proven inevitable for binding stability, structural integrity, and function (Table 1) (Figure 5A, left panel). Interestingly, the equivalent residue was not part of any cliques in the unbound form of the protein. Also, the equivalent residue Val175 of BLIPII in the TEM-1/BLIPII complex (PDB code: 1JTD) was not part of any inter-protein cliques. This residue was part of only an intra-protein clique within BLIPII (Figure 5A, right panel). Similarly, in CS2, SSH residue Cys53 of ricin/A9 complex Tyr91of ricin/A3C8 complexes were found in inter-protein and intra-protein cliques, whereas the equivalent residues were not part of any cliques in the unbound forms (PDB code: 1IFT, 6CWK, for ricin and A9 respectively) (Figures 5B left and right respectively). In CS3, the SSH residue Tyr13 is also part of both inter-protein and intra-protein cliques (Figure 5C). Especially, the other residues in the clique are part of domain 2 of 1L4-BP, while Tyr13 belongs to domain 1. This observation agrees with the PRS analysis in which the maximum effect of Tyr13 perturbations was observed in domain 2 residues. These results emphasize the importance of SSH in communication between the partner proteins in the complex as well as in the abridgement of interfacial and non-interfacial residues.

**Figure 5.**
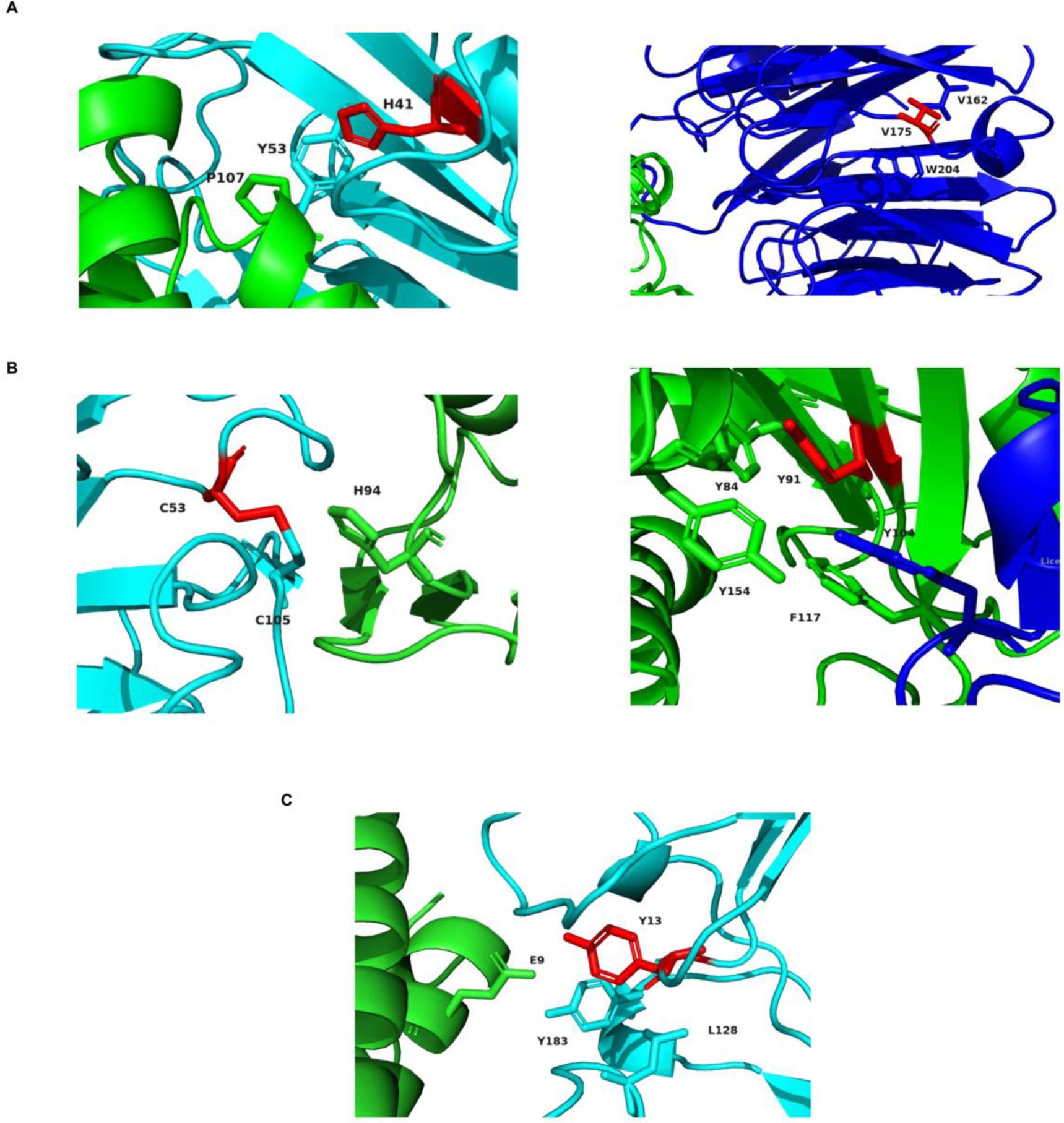
Cartoon representation from PyMol (The PyMOL Molecular Graphics System, Version 2.5 Schrodinger, LLC), with the residues that are part of cliques containing SSH in (A) TEM-1/BLIP complex (PDB code: 1JTG; left), TEM-1/BLIP1I complex (PD8 code: 1JTD; right panel), (B) Ricin/A9 complex (PDB code: 6CWG; left panel), ricln/A3C8 complex (PDB code: 5SV3; right panel), and (C) 1L-4/1L4-BP complex (PDB code: 11AR), shown in sticks. These cliques are predicted using Protein Sidechain Networks of each complex constructed using the web-server GraProStr61. TEM-1, ricin and IL-4 are shown in green cartoon, BLIP, A9, and IL4-BP in cyan cartoon and BLIPII and A3C8 in blue cartoon. The SSH in each case are shown in red sticks, and the other residues in the cliques are shown in sticks that are coloured according to their respective chains.

#### 3.5.7 Mutations of SSH are highly destabilizing

We performed *in silico* mutation of the SSH using the PositionScan module of FoldX^21^ to analyze how these mutations affect the binding and stability of the complexes (Figures S8A-D). When the SSH residue His41 of BLIP in CS1 was mutated to all other 19 amino acids, all the mutations led to a ΔΔG > 1 kcal/mol (Figure S8A left). However, when we mutated the equivalent residue Val175 of BLIPII in TEM-1/BLIPII complex (PDB code: 1JTD), Val, Ile, Pro had a ΔΔG < 1 kcal/mol (0, −0.003, and 0.6 kcal/mol respectively) (Figure S8A right). Similarly, when the SSH residue Cys53 of A9 in ricin/A9 complex (PDB code: 6CWG) of CS2 was mutated to any other 19 aa type, the destabilization was significant, as indicated by a ΔΔG > 3 kcal/mol, whereas the mutation of equivalent Arg53 in A3C8 which is not an SSH in ricin/A3C8 complex (PDB code: 5SV3) led to a ΔΔG > 1 kcal/mol only for four residue types, namely, Ser, Thr, Asn, and His (1.2, 1.2, 1.5, and 1.4 kcal/mol, respectively) (Figure S8B). The rest of the mutations had ΔΔG values between −1.9 kcal/mol and 1 kcal/mol. However, in the case of the other SSH Tyr91 of ricin, there were five mutants (Leu, Met, Phe, Tyr, and Trp) which had ΔΔG values in the range of −1 kcal/mol to 1 kcal/mol for both ricin/A9 and ricin/A3C8 complexes and which are neither stabilizing nor destabilizing in nature (Figure S8C left, and right panels respectively). Every other mutation leads to a destabilization. However, it is interesting to note that the position-specific mutations of an SSH and its topological equivalent conserved to be an SSH in the homologous complex have the same effect on both complexes. The result holds true in CS3 except for the Phe mutant of Tyr13 (0.3 kcal/mol), which has a ΔΔG between 0 kcal/mol and 1 kcal/mol (Figure S8D). Overall, these results suggest that mutating SSH to any residue type other than the wild type causes substantial destabilization of the complex, demonstrating the importance of these special hotspots.

## 4. CONCLUSIONS

The detailed analysis of SSH revealed their unique properties and biological roles in the protein-protein complexes. Compared to the ASA-hotspots and common-hotspots, SSHs tend to stay buried in both the bound and isolated forms of the protein. Hence, they were observed to have very distinct interaction patterns and a compact environment compared to the other hotspots. Compared to others, these hotspots are more evolutionarily conserved and prefer to be in structured secondary structures. There is a higher propensity for hydrophobic and long chain residues owing to their unique property of being less solvent accessible yet forming stabilizing interactions with the partner protein. The detailed case study analyses showed that the interaction network formed by the SSH is crucial for the activity and stability of the complex and remains conserved across homologous complexes. The PRS profiles showed that the SSHs are effectors whose perturbations have the most effect on non-interfacial sites. However, they are not sensors, but the perturbations on most interfacial sites have the highest impact on them. Protein sidechain network analyses showed that the SSH are also part of inter-protein and intra-protein cliques. Overall, the results suggest the importance of this special category of hotspots in the activity and stability of the complex, communication among the proteins in the complex, and bridging the interfacial and non-interfacial regions of proteins in the complex. Hence, this study suggests that the SSHs can be selected as restraints for various docking algorithms and as attractive medical targets for designing inhibitor drugs. We hope these findings help in improving the strategies for developing inhibitors targeting protein-protein interactions.

## Acknowledgements

This research is supported by Parvathy’s PMRF grant and NS’s JC Bose fellowship. Financial support for Parvathy Jayadevan from the Prime Minister’s Research Fellows (PMRF) scheme and the financial support for Yazhini Arangasamy from the Manfred Eigen fellowship from Max Planck Institute of Multidisciplinary Sciences is gratefully acknowledged. Srinivasan Narayanaswamy is a J.C. Bose National Fellow. Sowdhamini Ramanathan is a J.C. Bose National Fellow (JBR/2021/000006) from the Science and Engineering Research Board, India and Bioinformatics Centre Grant funded by the Department of Biotechnology, India (BT/PR40187/BTIS/137/9/2021). RS would also like to thank the Institute of Bioinformatics and Applied Biotechnology for the funding through her Mazumdar-Shaw Chair in Computational Biology (IBAB/MSCB/182/2022).

## Author Contributions

RS and NS conceptualized the study, and RS coordinated the study. PJ did all the analyses and wrote the first draft of the manuscript. PJ and YA reviewed the results, and YA and RS improved the manuscript.

## Data Availability Statement

PDB codes used in this study can be accessed from the RCSB Protein Data Bank.

