## Supplementary material for "Understanding the roles of secondary shell hotspots in protein-protein complexes": 27_8_24_biorxiv_supplementary.pdf

### Supplementary figures and tables

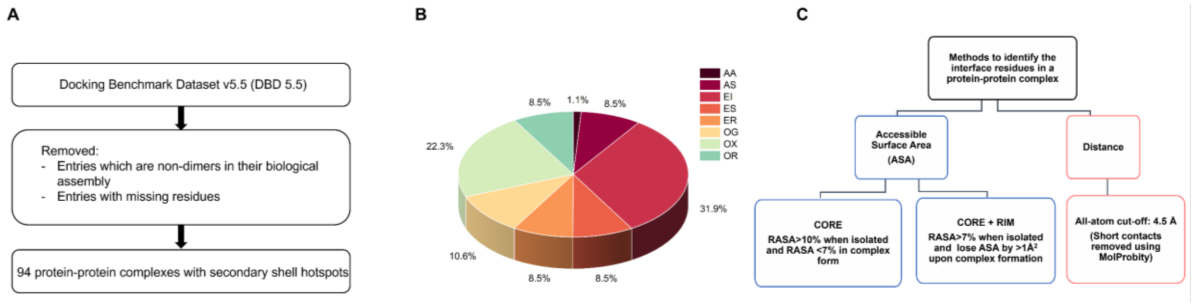


**Figure S1** (A) Flowchart of dataset generation. (B) Piechart showing the percentage of different types of protein-protein complexes in the dataset, namely enzyme-inhibitor (EI), enzyme-substrate (ES), enzyme complex with a regulatory or accessory chain (ER), antibody-antigen (AA), antigen-single domain antibody (AS), others, G-protein containing (OG), others, receptor containing (OR), and others, miscellaneous (OX), as taken from Docking Benchmark 5.5[1]. (C) Distance and ASA-based criteria that were used to identify interface residues in protein–protein complexes in the dataset.

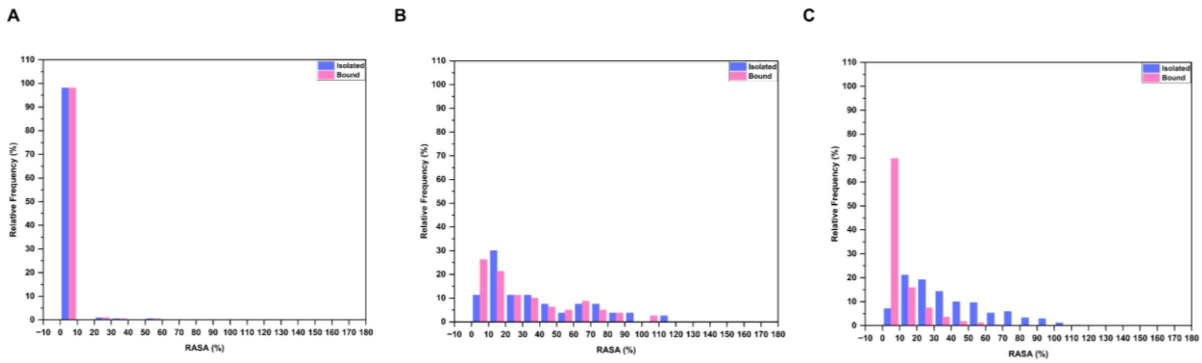


**Figure S2** Percentage of (A) SSH, (B) ASA-hotspots, and (C) common-hotspots with RASA values in bins of size 10% ranging from 0 to 180%, in the bound and isolated forms of the protein. RASA values were predicted using NACCESS[2].

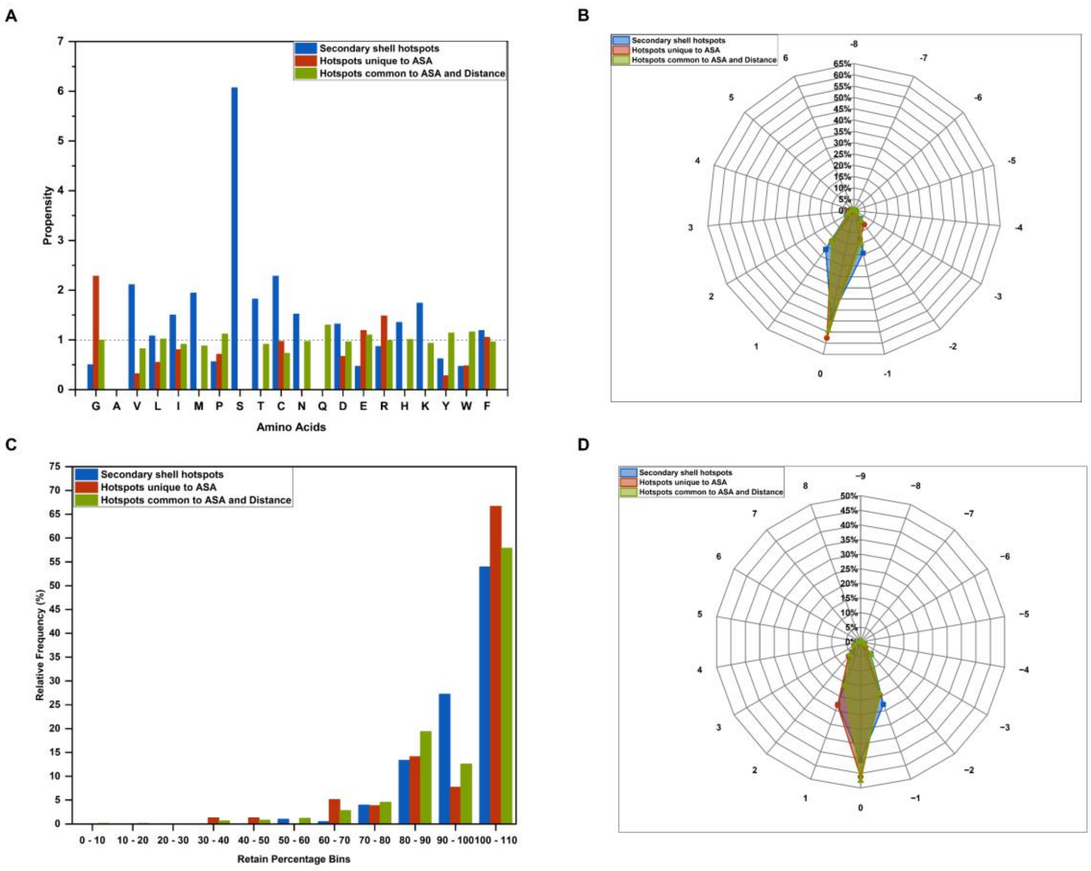


**Figure S3** (A) Chou-Fasman[3] propensities for each of the 20 amino acid types, for being SSH, ASA-hotspots, and common-hotspots. Plots showing the percentage of SSH, ASA-hotspots, and common-hotspots with (B) change in Ooi[4] number upon complexation varying from -8 to 6, and (C) retain percentage in bins of size 10%, ranging from 0% to 100%, and (D) change in the number of intra-protein partners from unbound to bound, varying from -9 to 8. Ooi number was predicted using SSTRUC (D. Smith, unpublished results), and intra-protein partners were identified using an in-house Python script that employs an all-atom distance-based criteria with a cut-off of 4.5 Å.

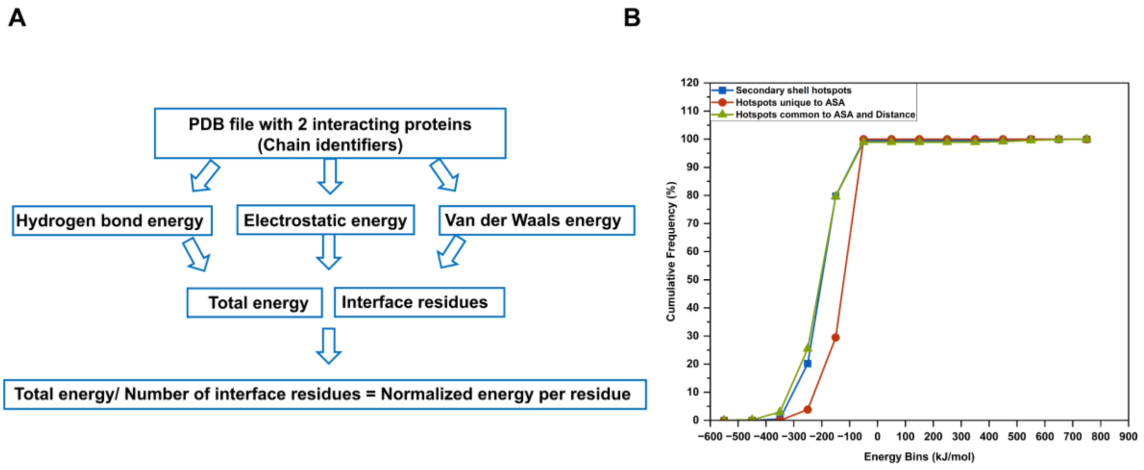

**Figure S4** (A) PPCheck[5] methodology. (B) Cumulative frequency in percentage of SSH, ASA-hotspots, and common-hotspots with residue-wise energy in bins of size 100 kJ/mol, ranging from -600 kJ/mol to 800 kJ/mol. Residue-wise energies were predicted using PPCheck.

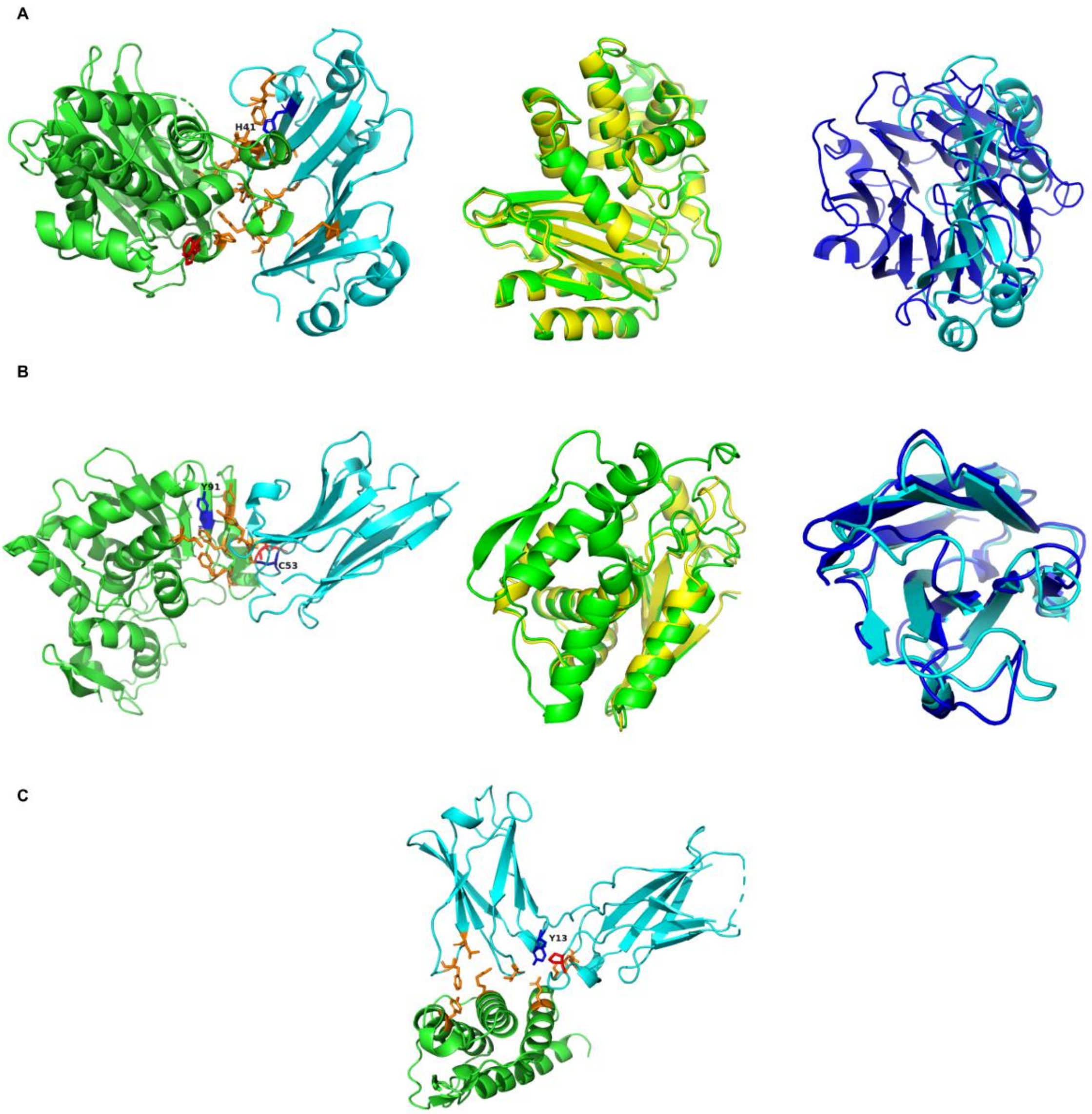


**Figure S5** Cartoon representation of (A) TEM-1/BLIP complex (PDB code: 1JTG; left panel), TM-align[6] superpositions of TEM-1 from TEM-1/BLIP complex (PDB code: 1JTG) and TEM-1/BLIPII complex (PDB code: 1JTD) (middle panel), and TM-align superpositions of BLIP from TEM-1/BLIP complex (PDB code: 1JTG) and BLIPII from TEM-1/BLIPII complex (PDB code: 1JTD) (right panel) of CS1, (B) ricin/A9 complex (PDB code: 6CWG; left panel), TM-align superpositions of ricin from ricin/A9 complex (PDB code: 6CWG) and ricin/A3C8 complex (PDB code: 5SV3) (middle panel), and TM-align superpositions of A9 from ricin/A9 complex (PDB code: 6CWG) and A3C8 from ricin/A3C8 complex (PDB code: 5SV3) (right panel), of CS2 and (C) IL-4/IL4-BP complex (PDB code: 1IAR) of CS3, generated using PyMol (The PyMOL Molecular Graphics System, Version 2.5 Schrödinger, LLC). SSH, ASA-hotspots and common-hotspots are shown in blue, red, and orange sticks. TEM-1 from TEM-

1/BLIP complex (PDB code: 1JTG), ricin from ricin/A9 complex (PDB code: 6CWG) and IL-4 from IL-4/IL4-BP complex (PDB code: 1IAR) are shown in green cartoon, TEM-1 from TEM-1/BLIPII complex (PDB code: 1JTD), ricin from ricin/A3C8 complex (PDB code: 5SV3) in yellow cartoons, BLIP, A9, and IL4-BP in cyan cartoon and BLIPII and A3C8 in blue cartoon.

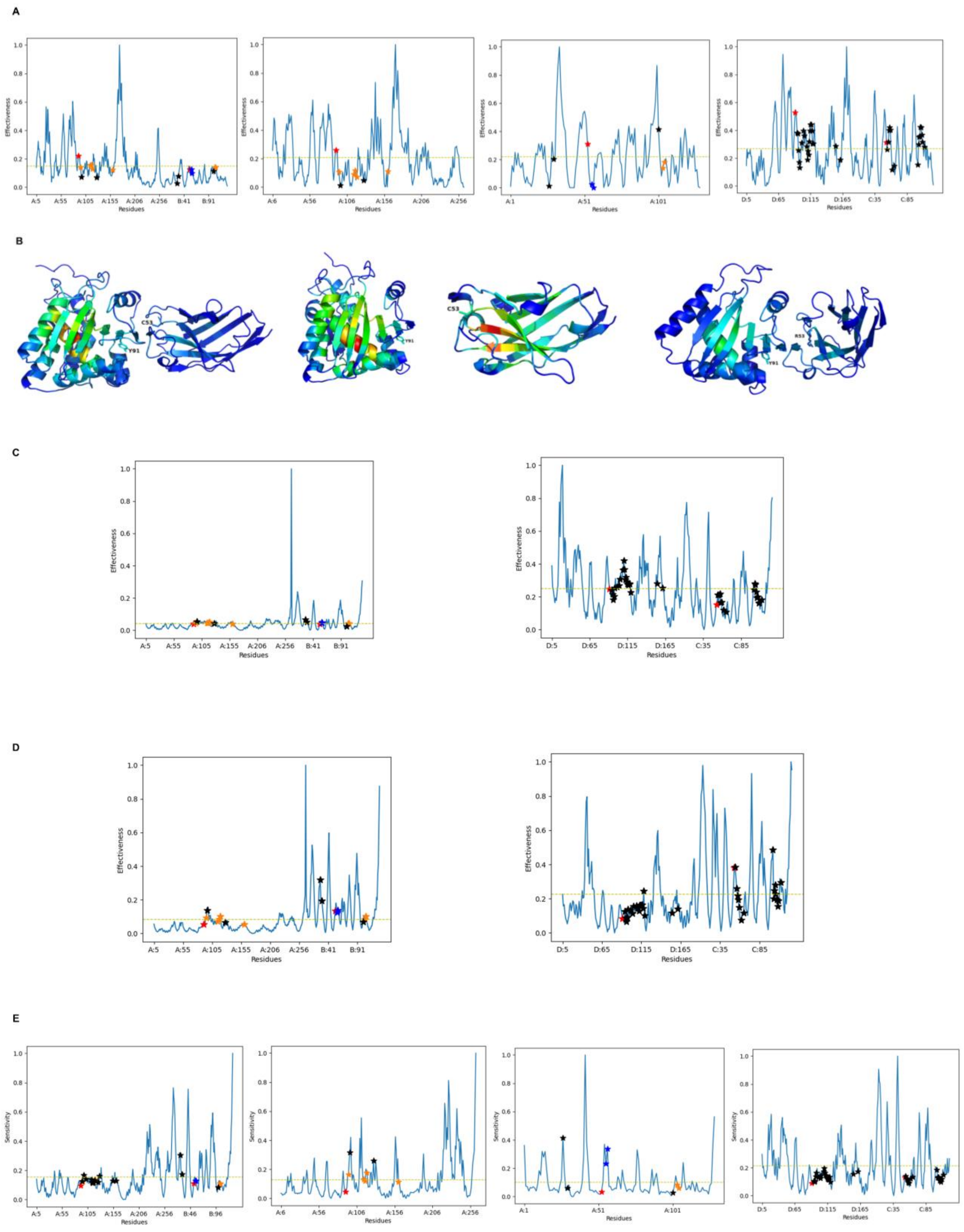

F

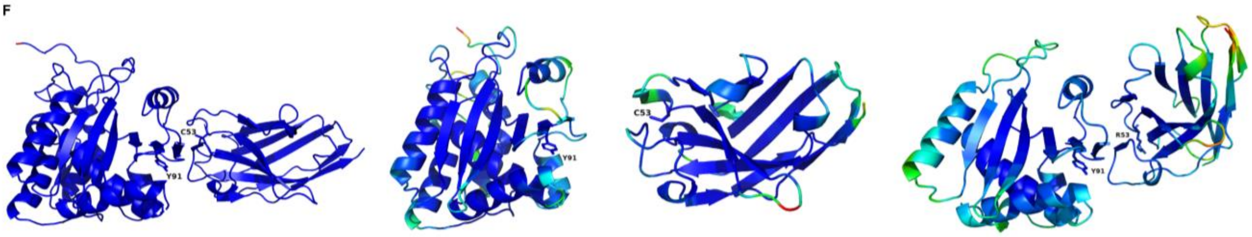


G

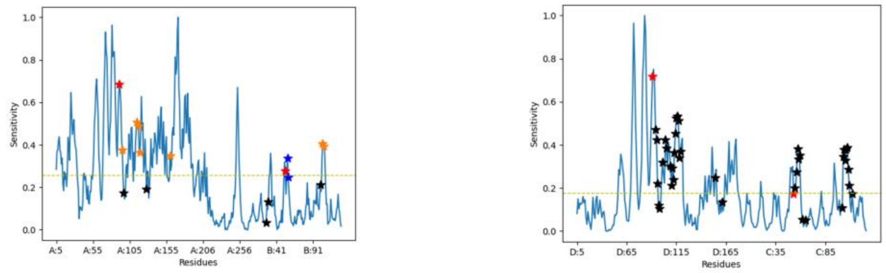


H

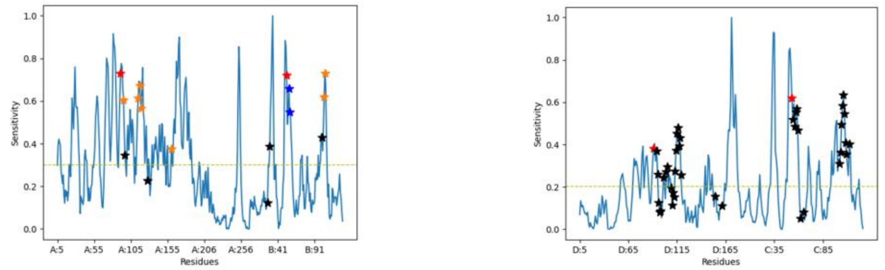


**Figure S6** (A) Normalized effectiveness profile and (B) Cartoon representation of ricin/A9 complex (PDB code: 6CWG; from left panel), unbound structure of ricin (PDB code: 1IFT), unbound structure of A9 (PDB code: 6CWK), and ricin/A3C8 complex (PDB code: 5SV3), with effectiveness values mapped (figure generated using PyMol, The PyMOL Molecular Graphics System, Version 2.5 Schrödinger, LLC). Residue-wise effectiveness values for all residues in ricin/A9 (PDB code: 6CWG; left panel) upon perturbing the SSH (C) Tyr91 of ricin, and (D) Cys53 of A9 and for all residues in ricin/A3C8 (PDB code: 5SV3; right panel), upon perturbing the equivalent residues Tyr91 of ricin and Arg53 of A3C8, respectively. (E) Normalized sensitivity profile and (F) cartoon representation of ricin/A9 complex (PDB code: 6CWG; from left panel), unbound structure of ricin (PDB code: 1IFT), unbound structure of A9 (PDB code: 6CWK), and ricin/A3C8 complex (PDB code: 5SV3), with sensitivity values mapped (figure generated using PyMol, The PyMOL Molecular Graphics System, Version 2.5 Schrödinger, LLC). Residue-wise sensitivity values of the SSH (G) Tyr91 of ricin, and (H) Cys53 of A9 upon perturbing all residues in ricin/A9 (PDB code: 6CWG; left panel), and the equivalent residues Tyr91 of A9 and Arg53 of A3C8, upon perturbing each residue in the ricin/A3C8 complex (PDB code: 5SV3; right panel). These profiles are generated by performing perturbation response scanning using the Prody[7] package. The effectiveness and sensitivity values in all profiles are normalized using the min-max method. The mean of effectiveness and sensitivity of all residues is considered as a cut-off in each profile, above which a residue is considered an effector or sensor (shown as yellow dashed line). The SSH,

ASA-hotspots, common-hotspots, and the interfacial residues excluding these hotspots are marked in red, blue, orange, and black stars, respectively.

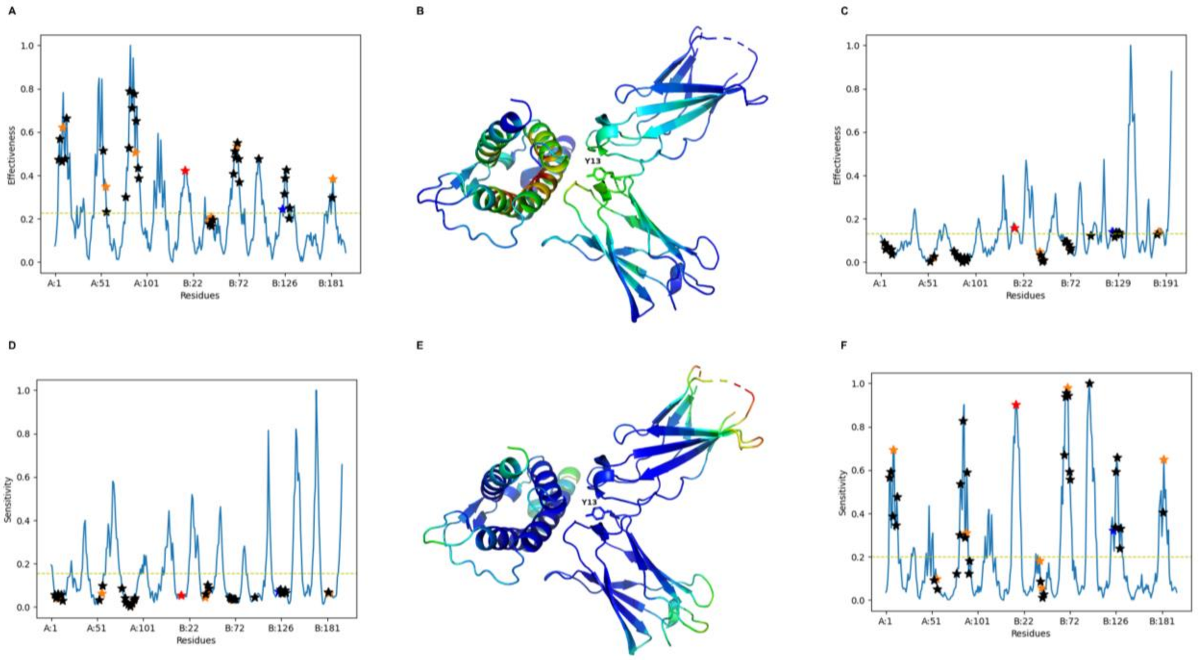


**Figure S7** (A) Normalized effectiveness profile and (B) Cartoon representation of IL-4/IL4-BP complex (PDB code: 1IAR), with effectiveness values mapped (figure generated using PyMol, The PyMOL Molecular Graphics System, Version 2.5 Schrödinger, LLC). (C) Residue-wise effectiveness values for all residues in IL-4/IL4-BP complex (PDB code: 1IAR) upon perturbing the SSH Tyr13 of IL4-BP. (D) Normalized sensitivity profile and (E) Cartoon representation of IL-4/IL4-BP complex (PDB code: 1IAR), with sensitivity values mapped (figure generated using PyMol, The PyMOL Molecular Graphics System, Version 2.5 Schrödinger, LLC). (F) Residue-wise sensitivity values of the SSH Tyr13 of IL4-BP upon perturbing each residue in the IL-4/IL4-BP complex (PDB code: 1IAR). These profiles are generated by performing perturbation response scanning using the Prody[7] package. The effectiveness and sensitivity values in all profiles are normalized using the min-max method. The mean of effectiveness and sensitivity of all residues is considered as a cut-off in each profile, above which a residue is considered an effector or sensor (shown as yellow dashed lines). The SSH, ASA-hotspots, common-hotspots, and the interfacial residues excluding these hotspots are marked in red, blue, orange, and black stars, respectively.

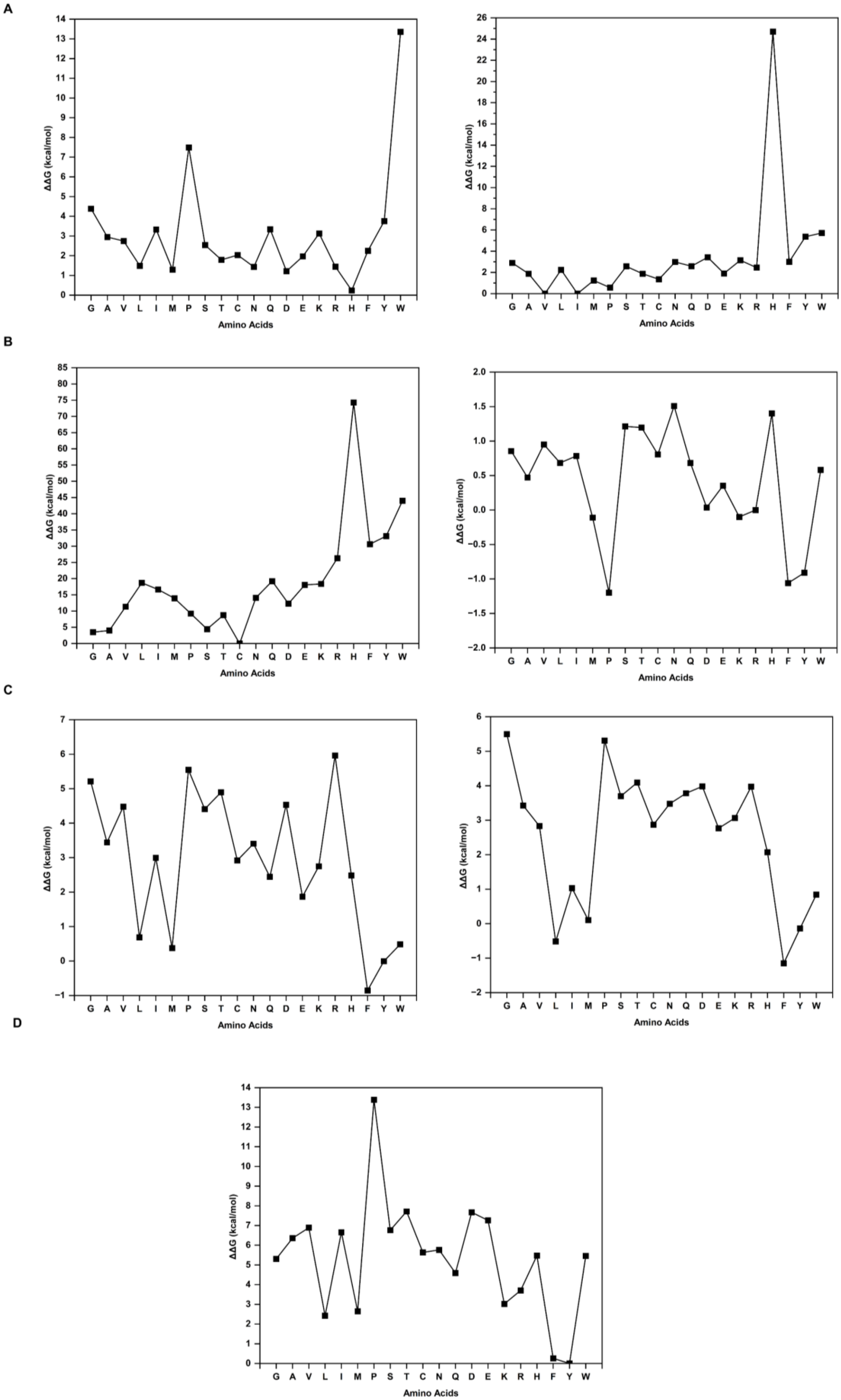
A
B
C
D
ΔΔG (kcal/mol)
Amino Acids
G A V L I M P S T C N Q D E K R H F Y W

**Figure S8** Line plots showing ΔΔG in kcal/mol upon position-specific *in silico* mutagenesis of SSH (A) His41 of BLIP in TEM-1/BLIP complex (PDB code: 1JTG; left panel), and the equivalent residue Val175 of BLIPII in TEM-1/BLIPII complex (PDB code: 1JTD; right panel), (B) Cys53 of A9 in ricin/A9 complex (PDB code: 6CWG; left panel), and the equivalent residue Arg53 of A3C8 in ricin/A3C8 complex (PDB code: 5SV3; right panel), (C) Tyr91 of ricin in ricin/A9 complex (PDB code: 6CWG; left panel), and the equivalent residue Tyr91 of ricin in ricin/A3C8 complex (PDB code: 5SV3; right panel), and (D) Tyr13 of IL4-BP in IL-4/IL4-BP complex (PDB code:1IAR), using PositionScan module of FoldX[8].

**Table S1:** RMSD[†] and sequence identity from TM-align[6] for CS1

| **Structure 1** | **Structure 2** | **RMSD from TM-align** | **Sequence ID from TM-align (%)** |
|---|---|---|---|
| TEM-1 from bound | TEM-1 from unbound | 0.3 | 100 |
| BLIP from bound | BLIP from unbound | 0.6 | 100 |
| BLIPII from bound | BLIPII from unbound | 0.3 | 100 |
| BLIP | BLIPII | 5.1 | 13.8 |

† RMSD: Root Mean Square Deviation, CS1: Case study 1

**Table S2:** RMSD[†] and sequence identity from TM-align[6] for CS2

| **Structure 1** | **Structure 2** | **RMSD from TM-align** | **Sequence ID from TM-align (%)** |
|---|---|---|---|
| Ricin from bound | Ricin from unbound | 0.5 | 100 |
| A9 from bound | A9 from unbound | 0.9 | 100 |
| A3C8 from bound | A3C8 from unbound | 0.6 | 100 |
| A9 from bound | A3C8 from bound | 1.8 | 67.7 |

† RMSD: Root Mean Square Deviation, CS2: Case study 2

**Table S3:** Interaction partners of the SSH Tyr91 of ricin in ricin/A9 complex (PDB code:6CWG) of CS2[†], the unbound structure of ricin (PDB code: 1IFT), and the equivalent residue Tyr91 of ricin in ricin/A3C8 complex (PDB code: 5SV3) of CS2. The inter-protein and intra-protein partners were identified using an all-atom distance cut-off of 4.5 Å, and the nature of interactions was predicted using the PIC[9] and RING[10] web-server. The importance of the residues in terms of structure, stability, and function was taken from earlier studies.

| **SSH in ricin/A9** | **Interaction partner in ricin/A9 bound** | | **Interaction partner in unbound** | **Importance** | **Equivalent in ricin/A3C8 bound** | **Interaction partner in ricin/A3C8 bound** |
|---|---|---|---|---|---|---|
| | **Inter-protein** | **Intra-protein** | **Intra-protein** | | | |
| Tyr91 of ricin | | | | | Tyr91 [SS]* | |

| | | | | | | |
|---|---|---|---|---|---|---|
| | Tyr104 [H‡,A‡] | | | A9's CDR3 forms a single hydrogen bond with α-helix D mediated by Tyr104 of A9 and Tyr154 of RTA, burying an additional 90 Å$^2$ with RTA[11]. | | Tyr104 [H,A] |
| | | Gly83 | Gly83 | | | Gly83 |
| | | Tyr84 [H,A,HB‡] | Tyr84 [H,A,HB] | | | Tyr84 [H,A,HB] |
| | | Ser89 | Ser89 | | | Ser89 |
| | | Ala90 | Ala90 | | | Ala90 |
| | | Phe92 | Phe92 | | | Phe92 |
| | | Phe93 | Phe93 | | | Phe93 |
| | | Tyr115 [H, A] | Tyr115 [H, A] | A common-hotspot | | Tyr115 [H,A] |
| | | | | Part of the major A9 binding site on RTA[11]. | | |
| | | Phe117 [H,A] | Phe117 [H,A] | A common-hotspot | | Phe117 [H,A] |
| | | | | Part of the major A9 binding site on RTA[11]. | | |
| | | Tyr154 [H,A] | Tyr154 [H,A] | Part of the major A9 binding site on RTA[11]. | | Tyr154 [H,A] |
| | | | | A9's CDR3 forms a single hydrogen bond with α-helix D mediated by Tyr104 of A9 and Tyr154 of RTA, burying an additional 90 Å$^2$ with RTA[11]. | | |
| | | Ser155 | Ser155 | A common-hotspot | | Ser155 |
| | | | | Part of the major A9 binding site on RTA[11]. | | |

†CS2: Case study 2

‡A: aromatic-aromatic, H: hydrophobic, and HB: H-bond

**Table S4:** Interaction partners of the SSH Cys53 of A9 in ricin/A9 complex (PDB code: 6CWG) of CS2†, the unbound structure of A9 (PDB code: 6CWK), and the equivalent residueArg53 of A3C8 in ricin/A3C8 complex (PDB code: 5SV3) of CS2. The inter-protein and intra-protein partners were identified using an all-atom distance cut-off of 4.5 Å, and the nature of interactions was predicted using the PIC[9] and RING[10] web-server. The importance of the residues in terms of structure, stability, and function was taken from earlier studies.

| **SSH in ricin/A9** | **Interaction partner in ricin/A9 bound** | | **Interaction partner in unbound** | **Importance** | **Equivalent in ricin/A3C8 bound** | **Interaction partner in ricin/A3C8 bound** |
|---|---|---|---|---|---|---|
| | **Inter-protein** | **Intra-protein** | **Intra-protein** | | | |
| Cys53 of A9 | | | | | Arg53 | |
| | His94 [HB‡] | | | | | NIL |
| | | Ser30 | Ser30 | | | Gly30 |
| | | Met31 | Met31 | | | Asp31 [I‡,HB] |
| | | Tyr32 | Tyr32 | | | Tyr32 |
| | | | Met33 | | | Gly33 |
| | | Ile51 | Ile51 | | | Ile51 |

| | | | | | | |
|---|---|---|---|---|---|---|
| | | Met52 | Met52 | | | Ser52 |
| | | Ser54 | Ser54 | | | Ser54 |
| | | Gly55 [HB] | Gly55 [HB] | Gly55-59 in CDR2: Removal of a single Gly residue (59) from A9's CDR2 significantly reduced binding affinity for RTA and eliminated toxin-neutralizing activity[11]. | | NIL |
| | | | | The crystal structure of A9 bound to RTA revealed that CDR2 contacts RTA's α-helix B. This interaction is noteworthy because mAbs and VHHs that bind α-helix B typically possess potent toxin-neutralizing activity. The primary interaction within this region occurs between A9 residue Gly55 and RTA residue Ala101[11]. | | |
| | | Gly56 [HB] | Gly56 [HB] | ASA-hotspot | | NIL |
| | | | | Gly55-59 in CDR2: Removal of a single Gly residue (59) from A9's CDR2 significantly reduced binding affinity for RTA and eliminated toxin-neutralizing activity[11]. | | |
| | | Gly57 | Gly57 [HB] | ASA-hotspot | | Thr55 [HB] |
| | | | | Gly55-59 in CDR2: Removal of a single Gly residue (59) from A9's CDR2 significantly reduced binding affinity for RTA and eliminated toxin-neutralizing activity[11]. | | |
| | | | | The tertiary structures of A9 in the free and RTA-bound forms were essentially identical. One notable deviation between these two structures occurred within the CDR2 region at Gly57[11]. | | |
| | | Gly58 [HB] | Gly58 [HB] | Gly55-59 in CDR2: Removal of a single Gly residue (59) from A9's CDR2 significantly reduced binding affinity for RTA and eliminated toxin-neutralizing activity[11]. | | |
| | | Arg74 | Arg74 | | | |
| | | Cys105 [D‡,HB] | Cys105 [D,HB] | Intraprotein-disulphide bridge with Cys53 linking CDR2 and 3 and part of β strand in CDR3[11]. | | |
| | | Ser106 | Ser106 | Part of β strand in CDR3 | | |

†CS2: Case study 2

‡HB: H-bond, I: ionic, D: disulphide bridge

**TABLE S5** Interaction partners of the SSH Tyr13 of 1L4-BP in IL-4/IL4-BP complex (PDB code: 1IAR) of CS3†. The inter-protein and intra-protein partners were identified using an all-atom distance cut-off of 4.5 Å, and the nature of interactions was predicted using the PIC[9] and RING[10] web-server. The importance of the residues in terms of structure, stability, and function was taken from earlier studies.

| **SSH** | **Interaction partner** | | **Importance** |
|---|---|---|---|
| | **Inter-protein** | **Intra-protein** | |
| Tyr13 of IL4-BP | | | |
| | Glu9 [HB‡] | | A common-hotspot |
| | | | Part of helix A (1-20), the region of major interface with 1L4-BP, and the putative γc- |

| | | | |
|---|---|---|---|
| | | | binding site, and show prominent structural alterations from unbound to bound[12]. |
| | | | Central to the first cluster, and major binding determinant. The high-resolution mutational analysis of IL-4 indicated that the side chain and charge of Glu9 provide the largest contribution to the stabilization of the complex. Part of the epitope that determines both association and dissociation[12,13]. |
| | | | Energetically coupled with the sidechain of SSH[14] |
| | | | E9A/Q: ΔΔG>3.5kcal/mol[13];<br>resulted in a loss in binding affinity by two to three orders of magnitude. |
| | | | E9K: increased the EC50 by more than 1000-fold[15]. |
| | | Ser11 [HB] | |
| | | Asp12 | |
| | | Met14 | M14A: same affinity as WT IL4-BP (no structural perturbations)[14] |
| | | Ser15 | S15A: same affinity as WT IL4-BP (no structural perturbations)[14] |
| | | Ile16 | |
| | | Val68 [H‡] | V68A: showed a decrease in binding affinity by 2.5-101-fold. Their effect on affinity is probably indirect, by destabilizing the active-site loop structures[14]. |
| | | Val69 | A common-hotspot |
| | | | Part of cluster 2 |
| | | | At the core of the IL-4/IL4-BP contact, but contribute only marginally to binding affinity[14]. |
| | | | V69A: 35 to 110-fold decreased affinity[14]. |
| | | Ser70 [HB] | Part of cluster 1 and interacts with Glu9, Thr6, Lys12, Thr13, Asn89 of IL-4. |
| | | Pro92 [H] | P92A: showed a decrease in binding affinity by 2.5-101-fold. Their effect on affinity is probably indirect, by destabilizing the active-site loop structures[14]. |
| | | Ser93 | Part of cluster 1. |
| | | Val96 [H] | Part of L4 loop that connects D1 and D2, but not used for ligand binding. |
| | | Asn126 [HB] | Between cluster 1 and 3 at the interface. |
| | | Tyr127 [H,A‡] | Part of cluster 1. |
| | | | Exhibits cooperative effects with the SSH. |
| | | | Y127A: 35 to 110-fold decreased affinity[14]. |
| | | | Y127F: binds IL-4 as strongly as wild-type IL4-BP. Thus, only the aromatic ring provides IL-4 binding affinity and not the hydroxyl group of Y127[14]. |
| | | Leu128 [H] | |
| | | Tyr183 [H, A] | A common-hotspot |
| | | | Major component of cluster 1 at the interface |
| | | | The hydroxyl group of Tyr183 forming an H-bond with IL-4 Glu9 in cluster 1 is the second main receptor binding determinant[14]. |
| | | | Exhibits cooperative effects with the SSH[14]. |
| | | | The binding of IL4-BP sidechain Y183 is coupled to all analyzed IL-4 sidechains in cluster I[14]. |
| | | | Y183A: ΔΔG=3.7; have a strongly reduced affinity that is more than 500-fold lower than that of wild-type IL4-BP. The minimal chemical alteration introduced in the Y183F variant make it very unlikely that the observed decrease in |

| | | | |
|---|---|---|---|
| | | | binding affnity originates from structural alterations[14]. |
| | | | Y183F: The affinity of this variant is only 2.5-fold lower than that of the Y183A variant, indicating that the hydroxyl group is responsible for the 210-fold decrease in affinity[14]. |

†CS3: Case study 3
‡HB: H-bond, H: hydrophobic, A: aromatic-aromatic
